# A Developmental Mechanism to Regulate Alternative Polyadenylation in an Adult Stem Cell Lineage

**DOI:** 10.1101/2024.03.18.585561

**Authors:** Lorenzo Gallicchio, Neuza R. Matias, Fabian Morales-Polanco, Iliana Nava, Sarah Stern, Yi Zeng, Margaret T. Fuller

## Abstract

Alternative Cleavage and Polyadenylation (APA) often results in production of mRNA isoforms with either longer or shorter 3’UTRs from the same genetic locus, potentially impacting mRNA translation, localization and stability. Developmentally regulated APA can thus make major contributions to cell-type-specific gene expression programs as cells differentiate. During *Drosophila* spermatogenesis, approximately 500 genes undergo APA when proliferating spermatogonia differentiate into spermatocytes, producing transcripts with shortened 3’ UTRs, leading to profound stage-specific changes in the proteins expressed. The molecular mechanisms that specify usage of upstream polyadenylation sites in spermatocytes are thus key to understanding the changes in cell state. Here, we show that upregulation of PCF11 and Cbc, the two components of Cleavage Factor II (CFII), orchestrates APA during *Drosophila* spermatogenesis. Knock down of *PCF11* or *cbc* in spermatocytes caused dysregulation of APA, with many transcripts normally cleaved at a proximal site in spermatocytes now cleaved at their distal site, as in spermatogonia. Forced overexpression of CFII components in spermatogonia switched cleavage of some transcripts to the proximal site normally used in spermatocytes. Our findings reveal a developmental mechanism where changes in expression of specific cleavage factors can direct cell-type-specific APA at selected genes.

## Introduction

Alternative cleavage and polyadenylation (APA) of mRNAs results from changes in the site at which the 3’ end cut that terminates nascent transcripts is made (Tian and Manley 2016; Gallicchio *et al*. 2023). For alternative polyadenylation sites located in the region of a gene encoding the 3’UTR, APA results in production of alternate mRNA isoforms that differ in 3’UTR length and content, with potentially profound effects on transcript fate, including translation, stability, and localization (An *et al*. 2008; Mayr and Bartel 2009; Mansfield and Keene 2012; Berry *et al*. 2022). More than 70% of human genes are subject to alternative polyadenylation, making APA a widespread biological phenomenon (Gruber and Zavolan 2019). The most intriguing APA instances are those where the choice of 3’ end cut site is developmentally regulated, leading, for example, to production of transcripts with longer 3’UTRs from specific genes in neurons or shorter 3’UTRs in differentiating male germ cells compared to precursor cells (Hilgers *et al*. 2012; Miura *et al*. 2013; Shan *et al*. 2017; Vallejos Baier *et al*. 2017; Grassi *et al*. 2019; Berry *et al*. 2022). Changes in 3’UTR length due to APA also occur in certain pathological states, such as the 3’UTR shortening of transcripts observed in some cancers (Mayr and Bartel 2009; Morris *et al*. 2012; Venkat *et al*. 2020). Because these changes in 3’UTR content can have dramatic effects on protein expression and thus cell state, cell-type-specific regulation of APA is emerging as an important developmental mechanism to promote robust cell fate decisions.

During cleavage and polyadenylation, the last step of mRNA synthesis, the nascent mRNA molecule is cleaved and a string of adenosines is added by nuclear PolyA polymerase to protect the newly made transcript from degradation and facilitate its export from the nucleus (Jing *et al*. 1999). Most mRNAs undergo cleavage and polyadenylation, with a notable exception being histone transcripts, which are not polyadenylated (Marzluff *et al*. 2008). Four well-conserved multi-protein complexes have been identified as components of the Cleavage and Polyadenylation machinery (Figure 1 A). The core complex CPSF (Cleavage and Polyadenylation Specificity Factor) is responsible for recognizing (via the CPSF30 and WDR33 subunits) the polyadenylation signal (PAS) and inducing cleavage of the nascent RNA molecule (via its CPSF73 subunit). The PAS is an hexamer (most commonly AAUAAA or a close variant) in the nascent mRNA usually ∼15-30 nucleotides upstream of the cleavage site (Clerici *et al*. 2018; Gallicchio *et al*. 2023). Another complex, CstF (Cleavage and Stimulation Factor), recognizes U/GU-rich regions downstream of the PAS (MacDonald *et al*. 1994). CFI (Cleavage Factor I) recognizes a UGUA motif upstream of the PAS (Yang *et al*. 2011) and CFII (Cleavage Factor II) preferentially binds to G-rich sequences via its PCF11 subunit (Schäfer *et al*. 2018). The Cleavage and Polyadenylation machinery is also associated with several additional proteins: Symplekin (a scaffolding protein), PABP2, and the nuclear PolyA polymerase (Figure 1 A) (Gallicchio *et al*. 2023). While the central role of the core complex CPSF in recognizing the PAS and cleaving the nascent mRNA molecule is relatively clear, the specific contributions of the other complexes and associated factors in specifying whether, where, on what specific gene products, and in which cell types a cut is made are still under investigation (Vallejos Baier *et al*. 2017; Zhu *et al*. 2018; Turner *et al*. 2020).

**Figure 1.**
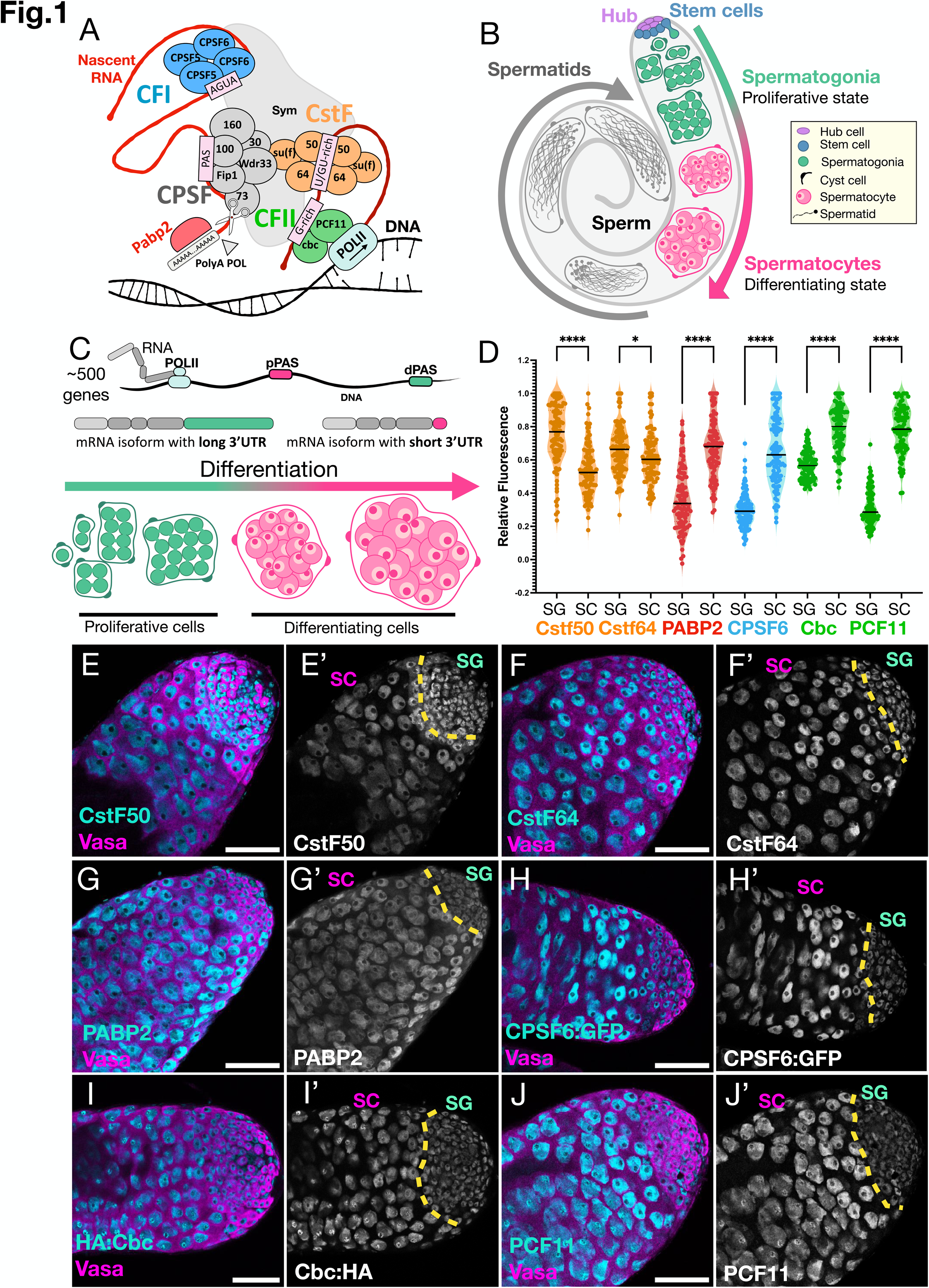
Expression of Cleavage Factors during *Drosophila* spermatogenesis. (A) Schematic representation of the cleavage machinery, adapted from Gallicchio *et al*. (2023). Grey: CPSF = Cleavage and Polyadenylation Factor complex. Orange: CstF = Cleavage and Stimulation Factor complex. Blue: CFI = Cleavage Factor I. Green: CFII = Cleavage Factor II. PAS = Polyadenylation signal. (B) Diagram of a wild-type *Drosophila* testes showing key stages of germ cell differentiation. (C) Diagram of developmentally regulated APA in the *Drosophila* testis resulting in expression of transcripts with long 3’UTRs in spermatogonia but short 3’UTRs in spermatocytes from ∼500 genes. (D) Quantification of relative immunofluorescence signal per nuclear area from (E-J). Each dot corresponds to a single nucleus. For each testis, 20 spermatogonial nuclei and 20 spermatocyte nuclei were quantified and N=5 testes were quantified per staining, for a total of 100 spermatogonia and 100 spermatocytes. Statistical significance calculated using the Kolmogorov-Smirnov t-test: (****) p-value < 0.0001; (*) p = 0.0366. (E-J) Example immunofluorescence images of testis apical regions from samples quantified in D. (Magenta) anti-Vasa marks germ cells. (Cyan) cleavage factor proteins: (E-G, J) Apical region of wild-type (*w1118*) testis stained with anti-(E) CstF50, (F) CstF64, (G) PABP2 and (J) PCF11. (H) Apical regions of testes from *CPSF6:GFP* transgenic fly stained with anti-GFP, (I) Apical region of testes from *HA:cbc/HA:cbc* fly stained with anti-HA. Dotted yellow line: border between proliferating spermatogonia (SG) and differentiating spermatocytes (SC). Scale bars = 50 uM.

Developmentally regulated changes in the PAS site utilized can have profound effects on the suite of proteins expressed as cells progress from one state to the next in a developmental sequence. For example, during spermatogenesis in flies and mice, many genes express mRNA isoforms with long 3’UTRs in proliferating spermatogonia, but isoforms with shorter 3’UTRs at later stages of germ cell development (Li *et al.,* 2016; Lee *et al.,* 2022; Berry *et al.,* 2022). In *Drosophila melanogaster* testes (Figure 1 B), over 500 genes express transcripts cleaved at a distal PAS in proliferating spermatogonia but at a more proximal PAS in differentiating spermatocytes, resulting in production of mRNA isoforms with much shorter 3’UTRs (Figure 1 C).

The switch from distal to proximal cleavage site usage occurs at a time of profound change in cell state in the male *Drosophila* germ line adult stem cell lineage. Spermatogenesis in *Drosophila* starts with an oriented stem cell division that causes one of the daughter cells to be displaced away from the somatic hub cells, stop self-renewing and start a program of transient amplification by mitotic proliferation, founding a clone of interconnected spermatogonia enclosed in two somatic cyst cells (Figure 1 B). After 4 rounds of mitosis the spermatogonia in the cyst start differentiation, beginning with pre meiotic S phase, producing 16 interconnected spermatocytes. After roughly 3.5 days of meiotic prophase, during which the germ cells grow 25 times in volume, the spermatocytes undergo meiosis I and II, producing a cyst of 64 haploid spermatids. These then undertake a series of morphological changes that lead to production of mature and functional sperm (Figure 1 B). The switch from spermatogonia to spermatocytes is accompanied by changes in both cell state and gene expression. Almost all the transcripts needed for the changes in cellular morphology that occur after meiosis are expressed in spermatocytes (Raz *et al*. 2023). Changes in protein expression due to 3’UTR shortening by APA may thus play key roles not only in setting up differences between proliferating spermatogonia vs differentiating spermatocytes, but also in stage-specific translational repression or activation required for proper timing of the subsequent cellular morphogenesis.

As the 3’UTR of mRNA molecules is a hub for *cis*-regulatory sequences that can attract *trans*-acting factors such as miRNAs and RNA binding proteins to regulate mRNA stability and translation efficiency, it is not surprising that the stage-specific 3’UTR shortening due to APA at over 500 genes observed in spermatocytes compared to spermatogonia (Figure 1 C) resulted in dramatic shifts in the proteins expressed between these two cell types. Polysome fractionation revealed that transcripts from at least 200 genes switched from the long 3’UTR isoform co-migrating with polysomes in spermatogonia to the short 3’UTR isoform not associated with ribosomes in young spermatocytes. For another ∼50 genes the long 3’UTR transcript isoforms were not translated in spermatogonia while the short 3’UTR isoforms co-migrated with polysomes in spermatocytes (Berry *et al.,* 2022). Thus, the developmental program(s) that specify whether nascent transcripts are cleaved at proximal 3’UTR sites in spermatocytes rather than the distal sites utilized in spermatogonia can switch expression of many proteins from ON in spermatogonia to OFF in spermatocytes, or vice versa, from OFF in spermatogonia to ON in spermatocytes.

Here we show that cell-type-specific production of transcripts with short 3’UTRs from specific genes due to APA in spermatocytes is developmentally regulated by changing the levels of one of the 4 complexes involved in 3’ end cleavage and polyadenylation. We found that the two protein components of CFII, encoded by the *PCF11* and *crowded by cid (cbc)* genes, are upregulated in spermatocytes compared to spermatogonia. When levels of *PCF11* or *cbc* were knocked down in spermatocytes by cell-type-specific RNAi, over 240 (∼48%) of the ∼500 genes for which nascent transcripts would normally be cleaved at a proximal PAS in spermatocytes instead resumed producing transcripts cleaved at the distal PAS utilized in spermatogonia. Reciprocally, forced overexpression of CFII components in spermatogonia shifted 3’ end cleavage to the more proximal site for some of the transcripts from genes that undergo stage-specific APA identified by Berry *et al*. (2022). Lowering protein levels in spermatocytes of components of the cleavage machinery associated complexes CFI or CstF had much less effect on the choice of 3’ end cleavage site, suggesting that CFII plays a special role in regulating APA in spermatocytes compared to spermatogonia. The PCF11 and Cbc proteins interact structurally: immunoprecipitation of Cbc from testis extracts brought with it a >250 KDa form of PCF11. Reduced expression of *PCF11* protein in spermatocytes due to RNAi resulted in concomitant reduction in Cbc, suggesting that Cbc protein was not stable without its binding partner in CFII. However, the reciprocal was not true. Plentiful PCF11 protein persisted in the nuclei of spermatocytes in which expression of *cbc* had been knocked down by RNAi. Strikingly, *PCF11* mRNA is broadly expressed in many cell types in adult flies, while *cbc* expression is low in most adult tissues but upregulated in spermatocytes. These observations suggest a model where a developmentally regulated increase in expression of *cbc* and *PCF11* in spermatocytes leads to greater activity of cleavage factor CFII. This in turn facilitates recognition of an upstream PAS and consequent 3’UTR shortening by APA of nascent transcripts from specific genes, indicating CFII as a key player in developmental regulation of alternative cleavage and polyadenylation.

## Results

### Specific cleavage factor proteins are upregulated in spermatocytes

Immunostaining of whole mount *Drosophila* testes revealed that some but not all cleavage factor proteins were upregulated in spermatocytes (Figure 1 D-J). Relative expression levels of six protein components of various sub-complexes of the cleavage factor machinery were measured by immunostaining using available antibodies against endogenous proteins (for CstF50, CstF64, PABP2 and PCF11), anti-HA (for *HA:cbc,* an available *Drosophila* line tagged with HA at the endogenous *cbc* locus by CRISPR (Wu *et al*. 2021)), or anti-GFP (for an available *Drosophila* line carrying a Fosmid based transgenic copy of CPSF6 tagged with sGFP (super folder GFP) (Sarov *et al*. 2016)). In comparing nuclear signal in spermatogonia and spermatocytes, each measurement was divided by the area of the corresponding nucleus to account for the great difference in nuclear size between the two cell types. Immunofluorescence signal from antibodies recognizing two members of the CstF complex, CstF50 (Figure 1 E) and CstF64 (Figure 1 F), decreased slightly or remained almost the same in nuclei in spermatocytes compared to nuclei in spermatogonia (quantified in Figure 1 D). In contrast, immunofluorescence signal from antibodies recognizing PABP2 (Figure 1 G), the CFI component CPSF6 (Figure 1 H) and the two members of CFII, Cbc (Figure 1 I) and PCF11 (Figure 1 J) increased substantially in nuclei from spermatocytes compared to nuclei from spermatogonia (quantified in Figure 1 D).

### Reduction of CFII components levels in spermatocytes shifted APA to more distal sites

To lower expression of cleavage factor components in spermatocytes, females carrying a *bamGal4* expression driver and *UAS>dicer2* were crossed with males carrying RNAi constructs under control of a *UAS* array with an *hsp70* promoter. The *bamGal4* flies express Gal4 under control of the *bag-of-marble (bam)* promoter, which fires in mid to late spermatogonial and early spermatocyte stages. The crosses were incubated at 25°C for three days, then the progeny shifted to 29°C to facilitate effective transcriptional activation by Gal4.

Immunofluorescence staining revealed that *bamGal4* driven knock down using the *VDRC-103710* RNAi line lowered levels of *PCF11* protein detected in spermatocyte nuclei to a level similar to that detected in spermatogonia (Figure 2 B). Testes from *UAS>PCF11-RNAi-VDRC-103710;bamGal4* males viewed in live squash preparations by phase contrast microscopy had plentiful spermatocytes but few elongating spermatid bundles (Figure 2 E). An RNAi line directed against a different region of the PCF11 transcript, *UAS*>PCF11-RNAi-*NIG-10228R-4*, also lowered the level of anti-PCF11 immunofluorescence signal in spermatocytes when driven using *bamGal4* (Figure 2 C). Testes from *UAS*>PCF11-RNAi-*NIG-10228R-4;bamGal4* males viewed by phase contrast microscopy had some elongating spermatid bundles (Figure 2 F). In both cases, PCF11 protein was still detected in early spermatogonia and in nuclei of the somatic cyst cells that accompany spermatocyte cysts, as expected since the *bamGal4* driver is not expressed in these cells (Figure 2 B, C, arrow heads).

**Figure 2.**
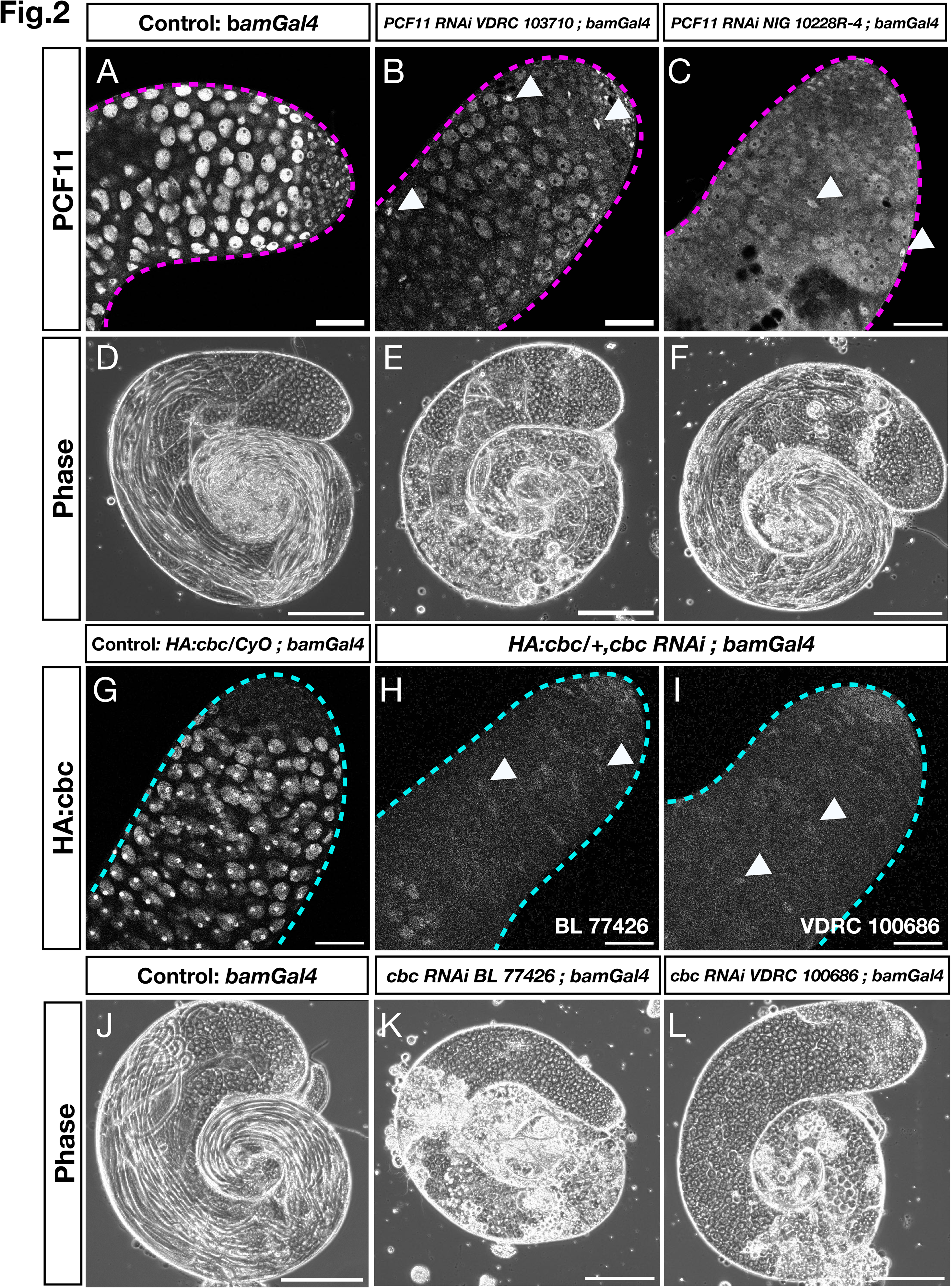
Knock down of *PCF11* and *cbc* in spermatocytes. (A-C) Apical tips of testes stained with anti-PCF11. (A) Control: *bamGal4*, (B) *PCF11-RNAi-VDRC-103710;bamGal4,* (C) *PCF11-RNAi NIG-10228R-4;bamGal4*. (D-E-F) Phase contrast microscope images of whole mount live testes from *Drosophila* lines in (A, B, C) respectively. (G-I) Apical tips of testes stained with anti-HA. (G) Control *HA:cbc/Cyo;bamGal4*, (H) *HA:cbc/+,cbc-RNAi-BL-77426;bamGal4* and (I) *HA:cbc/+,cbc-RNAi-VDRC-100686;bamGal4*. (J-L) Phase contrast microscope images of whole mount live testes. (J) Control: *bamGal4*, (K) *cbc-RNAi-BL-77426;bamGal4* and (L) *cbc-RNAi-VDRC-100686;bamGal4*. Arrow heads indicate somatic cyst cell nuclei. Scale bars are 50 uM (A-C, G-I) and 200 uM (D-F, J-L). Note that in all the immunofluorescence experiments on *HA:cbc/+* flies, the level of immunofluorescence signal with anti-HA in spermatogonia was below the detection limit for the experimental conditions. In contrast, for the testes quantified in Fig.1-I, which were from homozygous *HA:cbc*/*HA:cbc* flies, the level of HA:Cbc in nuclei of spermatogonia was high enough to be detected (for comparison between homozygous and heterozygous testes see SOM Figure 1).

Knock down of *cbc*, the partner of PCF11 in CFII, also caused defects in spermatogenesis. Immunofluorescence staining of testes from males heterozygous for a *cbc* allele tagged at the endogenous locus with the HA epitope (*HA:cbc*) showed strong anti-HA signal in spermatocyte nuclei, compared to little or no signal detected in spermatogonia. However, for two RNAi lines directed against different regions of *cbc*, knock down under control of *bamGal4* reduced the level of anti-HA signal in spermatocytes to below the level of detection, as in spermatogonia (Figure 2 G-I). In both cases, analysis of unfixed squash preparations by phase contrast microscopy showed that the knock down testes had abundant spermatocytes but lacked elongating spermatid bundles (Figure 2 K, L).

*In situ* hybridization revealed that knock down of *PCF11* or *cbc* in spermatocytes by RNAi affected APA at both the *Kpc2* (also named *isopeptidase-T-3*) and the *Red* genes (Figure 3). *Kpc2* and *Red* had both been found to express transcripts with a long 3’UTR in testis filled with spermatogonia but a much shorter 3’UTR in testes filled with mid-stage differentiating spermatocytes (Figure 3 A, E, J). The relative levels of transcripts with long vs. short 3’UTRs expressed in spermatogonia was compared by Hybridization Chain Reaction Fluorescence *in situ* Hybridization (HCR-FISH), a technique that can be used to quantify relative RNA levels in a sample, as the amplified fluorescent signal has been shown to be proportional to the number of mRNA molecules in the quantified area (Trivedi *et al*. 2018; Choi *et al*. 2018; Schulte *et al*. 2024). Comparing signal in the same sample from two different sets of HCR-FISH probes (SOM table 1), one recognizing the common coding sequence and one recognizing the 3’UTR extension (Figure 3 A), revealed the proportion of the total transcripts from a given gene (recognized by the common probe set) that contained the 3’ UTR extension (recognized by the 3’UTR extension probe). For both *Kpc2* and *Red*, knock down of either *PCF11* or *cbc* in spermatocytes under control of *bamGal4* substantially increased the level of signal from the 3’UTR extension detected in spermatocytes (Figure 3 C, D, H, I) compared to testes from control flies carrying the *bamGal4* driver but not the RNAi transgene raised under the same temperature regimen (Figure 3 B, G). Quantification of fluorescent signal confirmed that for both *Kpc2* and *Red* the ratio of fluorescence from probes recognizing the long 3’UTR versus probes against the coding sequence (CDS) for each gene was significantly higher in the *PCF11* or *cbc* knock down spermatocytes than in spermatocytes from control testes (Figure 3 F, K).

**Figure 3.**
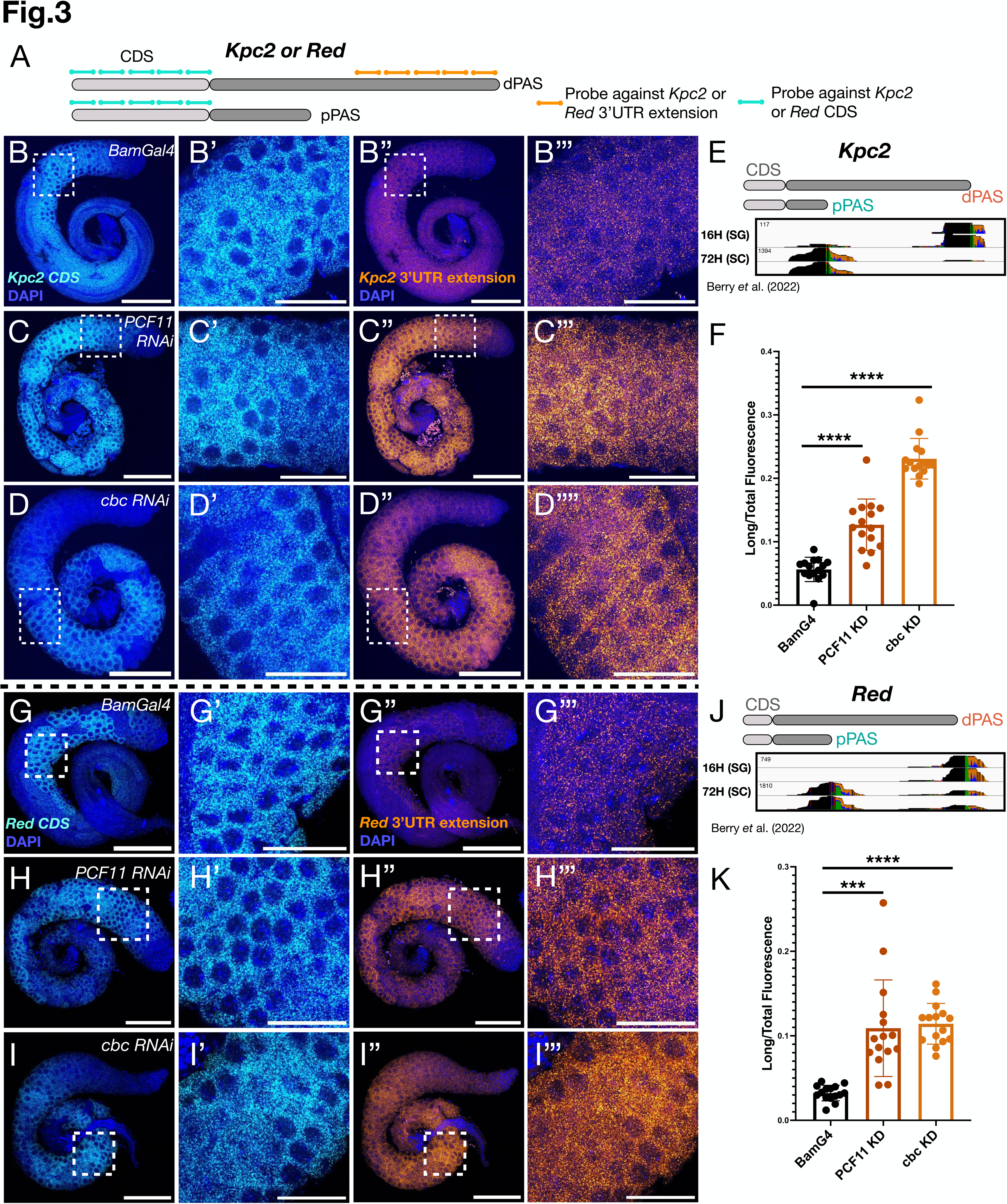
Knock down of either *PCF11* or *cbc* reduces cleavage at the proximal PAS in spermatocytes. (A) Diagram illustrating the design of the two probe sets. Light blue: probe against the protein coding sequence (CDS), orange: probe against the 3’UTR extension. (B-D’’) Confocal microscope images of (B, B’’) *bamGal4* (control), (C, C’’) *PCF11-RNAi-VDRC-103710;bamGal4* and (D, D’’) *cbc-RNAi-BL-77426;bamGal4* testes hybridized with probes recognizing the CDS of the *Kpc2* transcripts (cyan) and DAPI (B, C, D), and with probes recognizing the 3’UTR extension of the *Kpc2* transcripts (in orange) (B’’, C’’, D’’). (B’-D’, B’’’-D’’’) Zooms in spermatocytes of corresponding regions from (B-D, B’’-D’’) respectively. (E) IGV plots of 3’seq pile up of reads for the gene *Kpc2* from a sample enriched for SG = spermatogonia (16H PHS = post heat shock, Kim *et al*. 2017; Berry *et al*. 2022) and a sample enriched for spermatocytes (SC = spermatocytes, 72H PHS), to show the APA switch from the distal PAS (dPAS) used in spermatogonia and the proximal PAS (pPAS) used in spermatocytes. (F) Quantification of fluorescence ratio from probes recognizing the *Kpc2* 3’UTR extension (orange) compared to probes recognizing the *Kpc2* CDS (cyan) in spermatocytes. For each genotype N=3 testes were quantified. For each image 5 measurements were taken in different spermatocytes areas, which are plotted on the graph. (G-K) Analogous to (A-F) for the *Red gene*. Scale bars are 150 uM (B, B’’, C, C’’, D, D’’, G, G’’, H, H’’, I, I’’) and 50 uM (B’, B’’’, C’, C’’’, D’, D’’’, G’, G’’’, H’, H’’’, I’, I’’’). p values are (****) = p<0.0001 and (***) = p = 0.0001. For statistical analysis we used the Welch’s t-test.

Analysis of cleavage site usage by 3’seq from testes in which either *PCF11* or *cbc* had been knocked down in spermatocytes by RNAi under control of *bamGal4* showed that the increase in signal from the 3’UTR extension observed in knock down spermatocytes was due to a shift from cleavage at the proximal site normally utilized in spermatocytes to the more distal cut site normally utilized in spermatogonia for both *Kpc2* and *Red* (Figure 4 A, B). Knock down of *cbc* had the strongest effect. Knock down of *PCF11* shifted 3’ end cleavage to the same distal site as *cbc* knock down but to a lesser degree, especially with the *UAS>PCF11-RNAi-NIG-10228R-4* construct (Figure 3 A, B), consistent with the less efficient reduction in PCF11 protein levels observed with that particular line (Figure 2 C).

**Figure 4.**
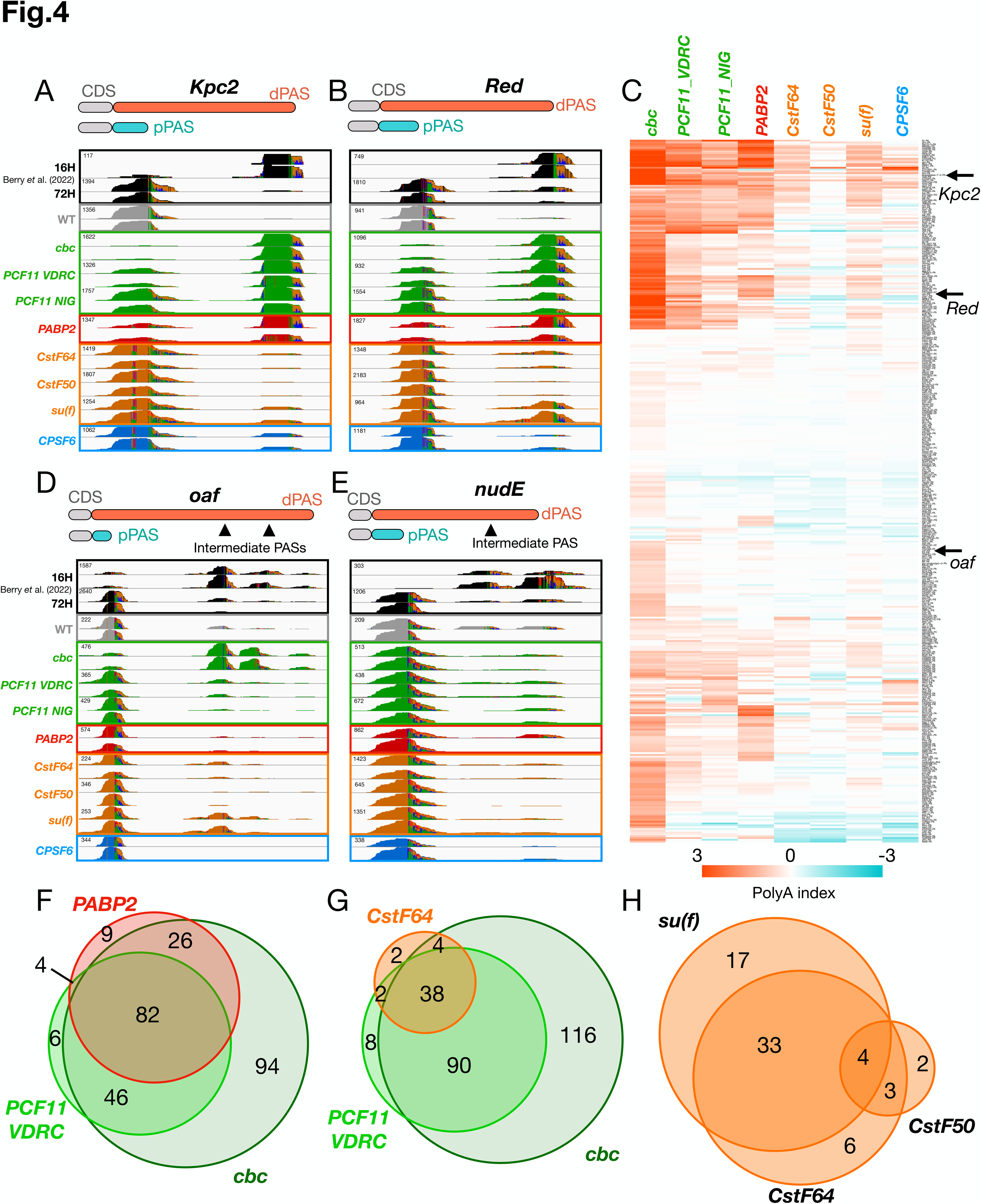
3’ end RNAseq comparing effects on APA of knock downs in spermatocytes of different components of the cleavage machinery. (A, B) Integrative Genomics Viewer (IGV) plots of 3’ end RNA sequencing reads for regions encoding the 3’UTR for the (A) *Kpc2* and (B) *Red* genes. Regions of peaks that change in color show mismatches between the PolyA tail and the genome to highlight the 3’ end of the mRNA. (C) Heatmap of PolyA index quantified with PolyA Miner across 3’seq libraries. Orange indicates increased cleavage at the distal PAS in knock downs compared to controls, and blue indicates increased cleavage at the proximal PAS in knock downs compared to control. Each row represents one of the >500 genes that switch to a proximal PAS in wild-type spermatocytes compared to spermatogonia from Berry *et al*. (2022). Each column represents a component of the cleavage machinery that was knocked down in spermatocytes. The interval chosen to plot the PolyA index (-3,3) does not represent the maximum and minimum PolyA index observed. In this heatmap genes that showed a statistically non-significant PolyA index in all the samples or a negative PolyA index in all the samples were removed, and in all other cases PolyA indexes that were non-significant were put equal to zero (another heatmap with these genes showed is reported in SOM Figure 2) (D, E) Integrative Genomics Viewer (IGV) plots of 3’ end RNA sequencing libraries for the genes (D) *oaf* and (E) *nudE*. (F-H) Venn diagrams representing number of genes with polyA index > 0.5 in *cbc* knock down libraries (dark green), *PCF11* VDRC knock down libraries (light green), *PABP2* knock down libraries (red) and components of CstF complex (orange).

3’seq data revealed that loss of function of *cbc* or *PCF11* affected APA at many loci. For roughly half of the ∼500 genes previously identified by Berry *et al*. (2022) as undergoing 3’UTR shortening by APA in spermatocytes compared to spermatogonia, knock down of either *PCF11* or *cbc* in spermatocytes resulted in increased usage of the distal cleavage site normally used in spermatogonia (Figure 4 C). When 3’seq data was analyzed via PolyA miner (Yalamanchili *et al*. 2020) providing as input the list of pairs of proximal PAS (pPAS) and distal PAS (dPAS) from the >500 genes undergoing APA identified by Berry *et al*. (2022), many genes showed increased usage of the distal PAS in knock down testes compared to control testes. Orange in Figure 4C indicates that the frequency of nascent transcripts from a given gene being cleaved at the distal PAS in knock down testes sample was higher than in control testes (positive PolyA index), while blue indicates that the frequency of the nascent transcripts being cleaved at the proximal PAS in knock down testes was higher than in control testes (negative PolyA index). Noticeably, a negative PolyA index does not indicate a shortening event but indicates a gene for which the proximal PAS was overrepresented in knock down compared to control samples. This could be caused either by different expression of the gene compared to in control testes or by the different cell composition in wild-type vs. knock down testes, as generally knock down testes do not have as many elongating spermatids as control testes. Again, knock down of *cbc* in spermatocytes had the strongest effect on the APA switch, affecting APA at the highest number of genes. Importantly, >90% of the genes where APA was affected by knock down of *PCF11* (PolyA index > 0.5, indicating more cleavage at the distal PAS) also showed a switch in APA to more cleavage at the distal site with knock down of *cbc,* consistent with these two CFII components acting together in the choice of where to make the 3’ end cut for transcripts from specific genes. Analysis of RNAseq libraries from testes samples in which *cbc* or *PCF11* were knocked down under *bamGal4* revealed that in general the genes that had a PolyA index either greater than 0.5 or smaller than negative 0.5 were not expressed at dramatically different levels in knock down versus control testes (SOM Figure 3).

3’seq revealed that lowering the levels of the nuclear PABP2 protein in spermatocytes by RNAi also frequently affected 3’UTR shortening by APA (Figure 4 A-C). Immunofluorescence staining showed effective knock down of PABP2 protein to below detection in spermatocytes from flies carrying *UAS>PABP2-RNAi-VDRC-106466;bamGal4*, although PABP2 protein was still abundant in nuclei of somatic cyst cells, as expected (SOM Figure 4 B). For two different RNAi lines, testes from *UAS>PABP2-RNAi;bamGal4* flies had plentiful spermatocytes but lacked elongating spermatid bundles (SOM Figure 4 D, E). 3’seq revealed that in testes from *UAS>PABP2-RNAi-VDRC-106466;bamGal4*, a substantial percentage of transcripts from the *Kpc2* and *Red* loci switched 3’ end cleavage from the proximal site normally utilized in spermatocytes to the more distal site normally utilized in spermatogonia (Figure 4 A, B). Analysis of the 3’seq data showed that for the >500 genes identified previously as undergoing APA in spermatocytes compared to spermatogonia,∼22% showed increased usage of the distal PAS when PABP2 was knocked down in spermatocytes. Furthermore, most (∼92%) of the genes that switched toward the more distal PAS (PolyA index > 0.5) after knocking down *PABP2*, also showed a shift (PolyA index > 0.5) towards the distal PAS when either *PCF11* or *cbc* was knocked down in spermatocytes (Figure 4 C, F).

In contrast, knock down of any of the three components of the CstF complex (*CstF64*, *CstF50*, and *su(f)*) or a component of CFI (*CPSF6*) in spermatocytes had much less effect on the APA profile of mRNAs from the ∼500 genes previously shown to undergo 3’ end shortening due to APA in spermatocytes, as assessed by 3’seq (Figure 4 C). Immunofluorescence staining with antibodies against CstF64 or CstF50 showed that levels of the proteins were substantially reduced in spermatocytes in testes in which expression of the respective genes had been knocked down by RNAi under control of *bamGal4* (SOM Figure 5). In both cases, signal remained in the nuclei of early spermatogonia and cyst cells, as expected from the cell-type-specificity of the *bamGal4* expression driver. No antibody reagents against *su(f)* were available to assess effectiveness of the RNAi construct directed against *su(f)*. However, the defects in spermatogenesis in *UAS>su(f)-RNAi;bamGal4* testes observed by phase contrast microscopy (SOM Figure 5) and the effects of knocking down *su(f)* on protein levels of CstF50 in spermatocytes nuclei (SOM Figure 6), suggested that the knock down of *su(f)* was effective. Levels of nuclear CstF50 protein drastically decreased in spermatocytes from males carrying an RNAi construct directed against *su(f)* expressed under control of *bamGal4* (SOM Figure 6 D). Knock down of *CstF64* under control of *bamGal4* also lowered the levels of CstF50 in spermatocyte nuclei (SOM Figure 6 F). Notably, when either *su(f)* or *CstF64* were knocked down in spermatocytes, nuclear CstF50 staining decreased, although some signal remained in the cytoplasm. Together the protein expression data suggest that components of the CstF complex may depend on each other for protein stability and/or nuclear localization, as previously reported in HeLa cells (Grozdanov *et al*. 2018). Consistent with the knock downs affecting function, squash preparations of testes from *UAS>su(f)-RNAI-BL-65693;bamGal4* showed abundant spermatocytes but no elongating spermatid stages (SOM Figure 5 G). Testes from *UAS>CstF64-RNAi-BL65987;bamGal4* males also showed defective spermatogenesis, with plentiful spermatocytes but extremely aberrant elongation stages (SOM Figure 5 F). Surprisingly, even though CstF50 protein was clearly strongly reduced in spermatocytes from *UAS>CstF50-RNAi-BL77377;bamGal4* males, spermatogenesis appeared largely normal by phase contrast microscopy (SOM Figure 5 C).

**Figure 5.**
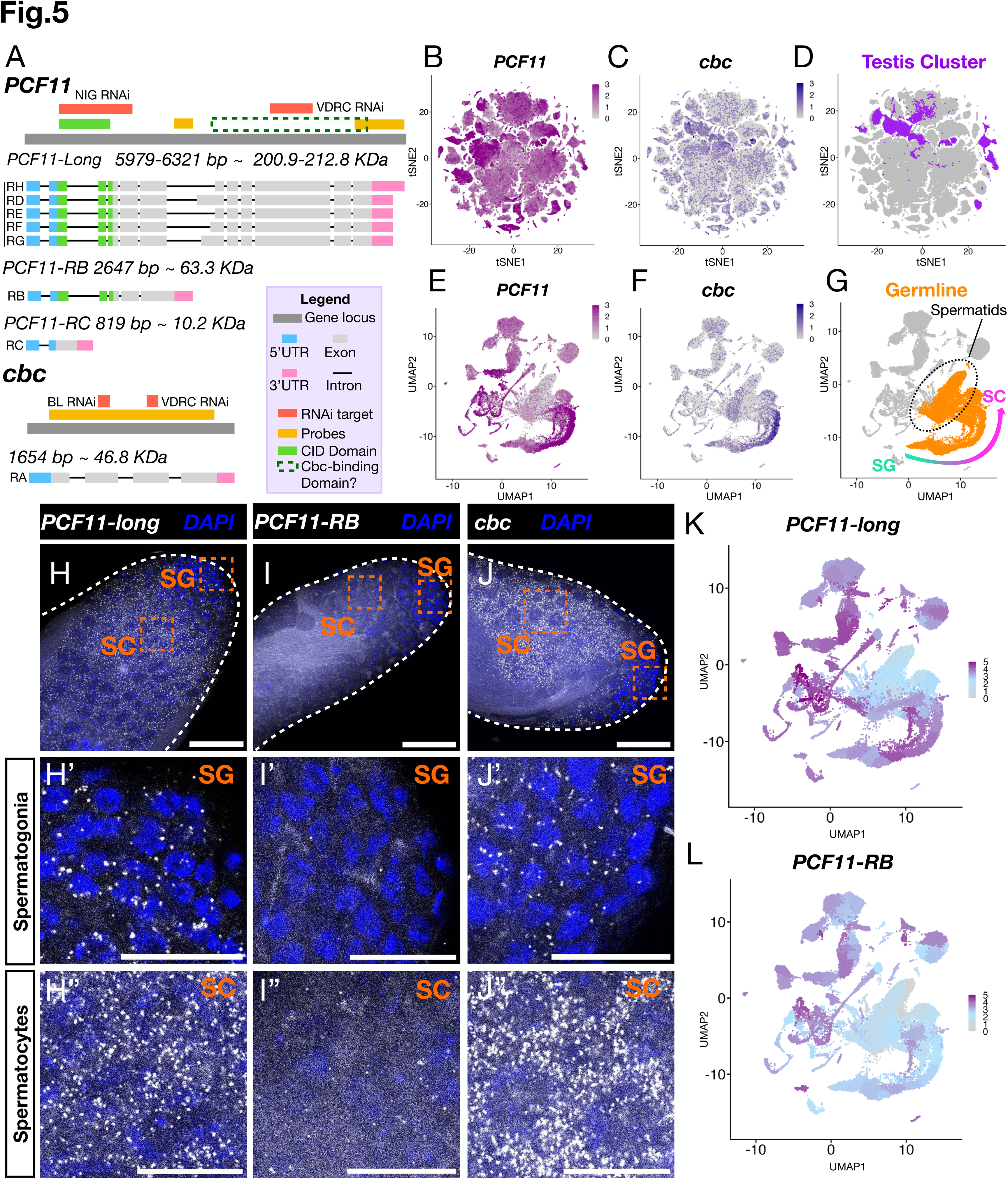
*PCF11* and *cbc* transcripts are both upregulated in spermatocytes with *cbc* particularly highly expressed in testis compared to other tissues. (A) Diagram adapted from FlyBase of *PCF11* and *cbc* loci. Light blue boxes represent 5’UTRs, green and grey boxes represent exons and pink boxes represent 3’UTRs. Introns represented with a straight black line. In yellow are represented the transcripts regions targeted by HCR-FISH probes for RNA *in situs*. Green and dotted-dark green boxes above the *PCF11* locus represent respectively the CID domain and a region in which the Cbc binding domain could be located. (B-G) Scope diagrams of Fly Cell Atlas single nuclear RNAseq data (Li *et al*. 2023) showing: (B) *PCF11* and (C) *cbc* expression across all the analyzed adult tissues, (D) Purple marking nuclei from the testis plus seminal vesicle sample across all analyzed adult tissues, (E-G) Umap plots showing (E) *PCF11* and (F) *cbc* expression in the testis plus semnal vesicle sample, (G) orange marking nuclei from the male germ-line clusters. (H-J) Confocal microscope images of apical tips of wild-type (*w1118*) testes hybridized with (H) probes against *PCF11* long isoforms (white) and DAPI (blue), (I) probes against *PCF11*-RB isoform (white) and DAPI (blue), (J) probes against *cbc* (white) and DAPI (blue). In (H, I, J) the boxes highlight a region of spermatogonia (SG) zoomed in (H’, I’, J’) and a region of spermatocytes (SC) zoomed in (H’’, I’’, J’’). (K, L) Umap from Fly Cell Atlas single nuclear data from testis plus seminal vesicle sample showing *PCF11-long* (K) and *PCF11-RB* (L) expression across the different clusters. Scalebars are 50 uM in (H, I, J) and 20 uM in (H’, H’’, I’, I’’, J’, J’’).

**Figure 6.**
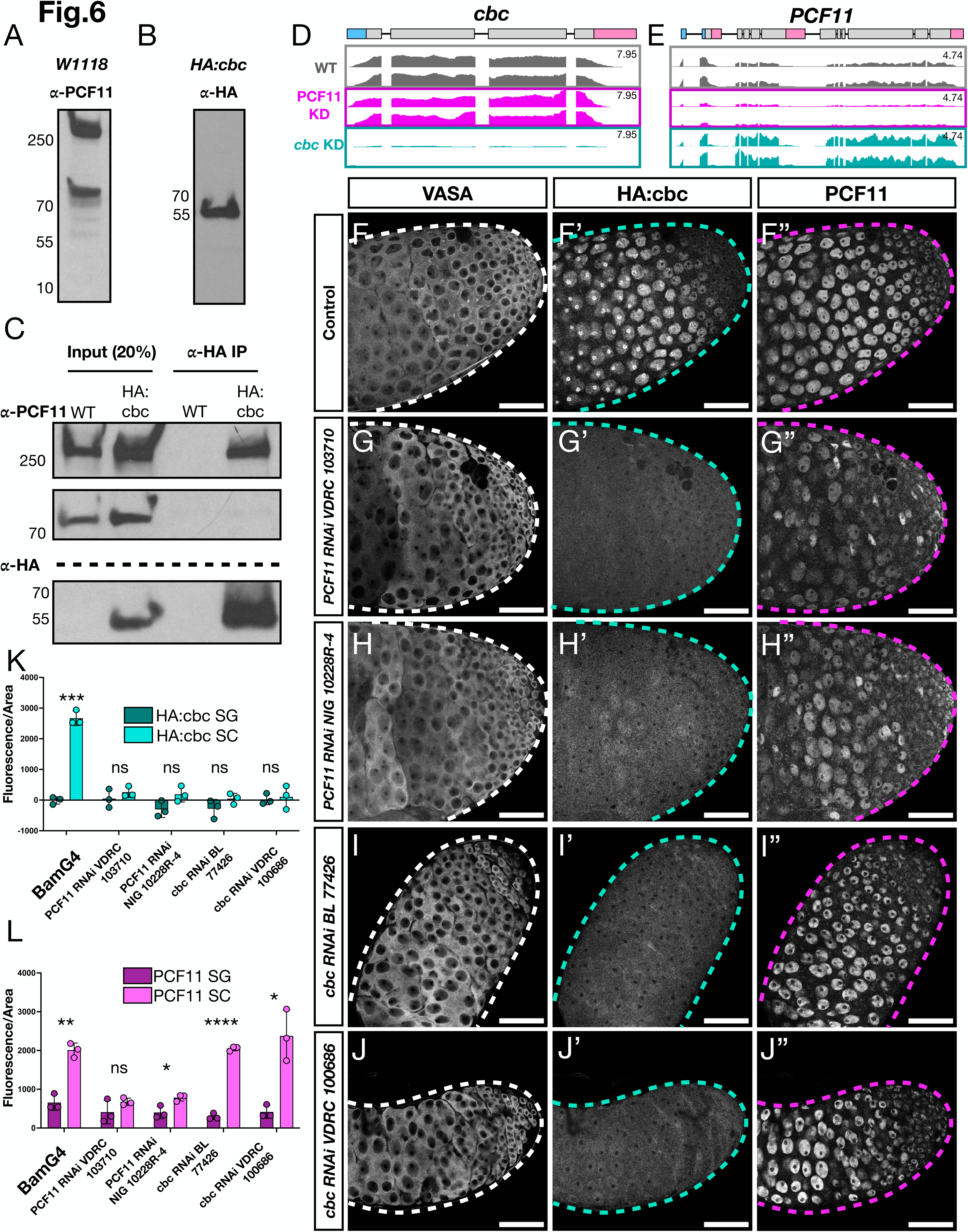
knock down of *PCF11* caused decrease of Cbc protein. (A) Western Blots of wild-type (*w1118*) testis lysate probes with anti-PCF11. (B) Western Blots of *HA:cbc* homozygous testes lysate probed with anti-HA. (C) Immunoprecipitation (IP) of *HA:cbc* in wild-type (*w1118*) and *HA:cbc* homozygous testes followed by PAGE and western blots probed with anti-PCF11 or anti-HA. Input = 20% of lysate before IP. (D, E) IGV plots of (D) *cbc* and (E) *PCF11* loci of RNAseq data from *bamGal4* (control) samples (in grey), *PCF11-RNAi-VDRC-103710;bamGal4* samples (magenta) and *cbc-RNAi-BL-77426;bamGal4* samples (teal). (F-J) Immunofluorescence images showing apical tips of testes stained with anti-Vasa to mark germ cells, anti-HA, and anti-PCF11. (F-F’’) *HA:cbc/CyO;bamGal4* (control), (G-G’’) *HA:cbc/+,PCF11-RNAi-VDRC-103710;bamGal4*, (H-H’’) *HA:cbc/+,PCF11-RNAi-NIG-10228R-4;bamGal4*, (I-I’’) *HA:cbc/+,cbc-RNAi-BL-77426;bamGal4* and (J-J’’) *HA:cbc/+,cbc-RNAi-VDRC-100686;bamGal4.* (K, L) Quantification of Fluorescence/Area from (K) HA antibodies (teal) and (L) PCF11 antibodies (magenta). For each testis 10 spermatogonial and 10 spermatocyte nuclei were quantified for both anti-PCF11 and anti-HA fluorescence and then averaged. For each genotype N=3 testes were quantified. Statistical analysis performed using the Welch’s t-test. p-values: *bamGal4_PCF11* = 0.0011 (**), b*amGal4_cbc* = 0.0005 (***), *PCF11-RNAi-NIG10228R-4_PCF11* = 0.0341 (*), *cbc-RNAi-BL77427_PCF11* < 0.0001 (****), *cbc-RNAi-VDRC100686_PCF11* = 0.0303 (*). Scalebars are 50 uM (F-J’’). Entire gels for western blots and co-IP are reported in SOM Figure 11.

Knock down of either *CstF64* or *su(f)* in spermatocytes shifted the site of 3’ end formation toward the more distal site utilized in spermatogonia at some of the >500 genes previously identified as subject to 3’UTR shortening in spermatocytes due to developmentally regulated APA (Figure 4 C). These genes tended to be among the same set of genes where APA was affected by knock down of components of CFII: ∼95% of the genes where APA was affected by knock down of *CstF64* were also affected by knock down of either *PCF11, cbc,* or both (Figure 4 G). However, the effects of loss of function of CstF components on APA in spermatocytes was small compared to the effect of loss of CFII components: only 44 of the 258 genes where APA was strongly affected by knock down of either *cbc* or *PCF11* in spermatocytes also showed a switch toward the more distal cut site in testes where function of *CstF64* had been knocked down in spermatocytes (Figure 4 G). More than 85% of the genes affected (PolyA index > 0.5) by *CstF64* knock down were also affected either by *CstF50* or *su(f)* knock down, and 68% of the genes affected by *su(f)* knock down were also affected by either *CstF64* or *CstF50* knock down (Figure 4 H), consistent with these three components acting together. For *CPSF6*, a component of the CFI complex, which binds nascent transcripts upstream the PAS, knock down under the control of *bamGal4* caused extensive defects in spermatid differentiation for one RNAi line (VDRC 107147) and variable or weaker defects for the other RNAi line tested (BL 34804, used for 3’seq), suggesting partial loss of function (SOM Figure 7). Analysis of cut sites utilized by 3’ end seq indicated that for the >500 genes previously identified as undergoing APA in spermatocytes, the effect of *CPSF6* knock down on APA using the BL 34804 RNAi line was weak compared to knock downs of *cbc* or *PCF11* (Figure 4 C). However, the weak effect could in part be due to a weaker knock down.

*De novo* genome wide calling of PAS sites located on annotated 3’UTRs revealed that knock down of components of the cleavage machinery affected APA at additional loci (SOM Figure 8 and SOM Figure 9) in addition to the >500 genes from Berry *et al*. (2022). For this analysis we only considered genes in which the most upregulated PAS and the most downregulated PAS in a knock down compared to a wild-type sample were located in the annotated gene 3’UTR. Again, the strongest effects were detected in *cbc* knock down testes, followed by *PCF11* knock down and *PABP2* knock down testes (SOM Figure 8), while *CstF64*, *CstF50*, *su(f)* and *CPSF6* knock downs had much milder effects on the global APA profile (SOM Figure 9).

Of the components of the cleavage machinery tested, the 3’seq data indicated that the 3’UTR shortening of transcripts from selected genes due to APA in spermatocytes was most strongly dependent on levels of CFII (composed of Cbc and PCF11), or *PABP2.* For almost half of the >500 genes previously identified as undergoing the developmentally regulated APA, lowering function of one or the other of these proteins in spermatocytes resulted in a shift toward usage of the more distal 3’ end cut site typical of spermatogonia. Several genes seemed to be more affected by knock down of *cbc* than by knock down of *PCF11*, for example *oaf* (Figure 4 D). Notably, over half of the genes previously identified as subject to 3’UTR shortening in spermatocytes compared to spermatogonia due to APA were not substantially affected by knock down of any of the cleavage factor machinery we tested, see *nudE* for example (Figure 4 E and SOM Figure 2).

### Transcription of the CFII components *PCF11* and *cbc* are strongly upregulated in spermatocytes

According to Fly Base, the *PCF11* locus can give rise to 7 different RNA isoforms (Figure 5 A), grouped in 3 classes for simplicity: a short isoform (*PCF11-RC*) of 819 nucleotides predicted to encode a small protein (∼10 KDa), a medium isoform (*PCF11-RB*) of 2647 nucleotides encoding a ∼63 KDa predicted protein, and a group of ∼6000 nucleotide transcripts (*PCF11-RD-RE-RF-RG-RH*) predicted to encode a family of >200 KDa proteins that have subtle differences in a central region. All the transcript isoforms share the same 5’UTR but differ in their 3’UTR (as well as in exon combinations). PCF11 has a CID domain (POLII C-terminal binding domain) present in both the PCF11-long and the 63 KDa PCF11-RB isoforms, but only partially present in the short protein encoded by *PCF11-RC*. PCF11 homologs in mammals and yeast have been shown to bind Cbc homologs through a region close to the C-terminus of PCF11 (Sadowski *et al*. 2003; Schäfer *et al*. 2018). Notably, while the *PCF11-RNAi-NIG10228-R4* line targets a region common to all *PCF11* mRNA isoforms, *PCF11-RNAi-VDRC107310* targets a region present only in the *PCF11-long* isoforms. The *cbc* locus is simpler, with only one predicted transcript 1654 nucleotides long, encoding a 46.8 KDa predicted protein (Figure 5 A).

The two *Drosophila* components of CFII show substantially different patterns of transcript expression in adult flies. In the Fly Cell Atlas single nuclear RNA-seq analysis of tissues from adult flies (Li *et al.,* 2022), transcription of *PCF11* was detected at generally high levels in many cell types (Figure 5 B). In contrast, *cbc* expression was detected at substantial levels in only a small subset of nuclei from adult tissues, with the clusters showing the highest *cbc* transcription belonging to the testes plus seminal vesicle sample (Figure 5 C, D). Plotting expression on the UMAP of snRNA-Seq data from testis plus seminal vesicle revealed that *PCF11* is transcribed in many somatic cell types in the testis, including terminal epithelia, seminal vesicle, nuclei from testis muscle cells and prominently in the somatic cyst cells that enclose germ line cysts. *PCF11* transcripts were also expressed in germ cells, with the level increasing substantially in spermatocytes compared to spermatogonia (Figure 5 E). Transcription of *cbc* also increased substantially in spermatocytes compared to spermatogonia (Figure 5 F).

Analysis of RNA expression in wild-type testes by HCR-FISH revealed that the long transcripts from *PCF11* were upregulated in spermatocytes compared to spermatogonia (Fig. 5 H-H’-H’’). In contrast, the intermediate sized transcript isoform *PCF11-RB*, detected with a set of probes recognizing the 3’UTR of *PCF11-RB*, which derives from an intronic region of the *PCF11-long* isoforms so should not recognize fully processed *PCF11-long* isoforms in the cytoplasm, was extremely low in both spermatogonia and spermatocytes (Figure 5 I-I’’). Single nuclear RNAseq from the Fly cell Atlas confirmed that while the collective *PCF11-long* isoforms are highly expressed in spermatocytes, cyst cells and somatic cell nuclei in the testis (Figure 5 K), the *PCF11-RB* transcript was only lowly expressed in the germline, with higher but still modest expression in cyst cell and other somatic cell nuclei in the testis (Figure 5 L). Similar HCR-FISH using a set of probes against the *cbc* protein coding region confirmed that expression of *cbc* mRNA was strongly upregulated in spermatocytes compared to spermatogonia (Figure 5 J-J’’).

### PCF11 and Cbc co-immunoprecipitate and knock down of *PCF11* causes decrease of Cbc protein in spermatocyte nuclei

Analysis of proteins expressed in wild-type testes by polyacrylamide gel electrophoresis (PAGE) followed by Western blot probed with antibodies against PCF11 showed a predominant high molecular weight (>250 KDa) band (Figure 6 A), consistent with the expression of *PCF11-long* mRNA isoforms detected by HCR-FISH and snRNA-seq (Figure 5 H, K). Anti-PCF11 also detected a ∼70 KDa protein isoform in testis extracts, as predicted from the *PCF11-RB* mRNA isoform. Although the antibody used was raised against amino acids 1-283 of *Drosophila* PCF11 (Zhang and Gilmour 2006) no protein product corresponding to PCF11-RC was observed, despite a seemingly high level of the *PCF11-RC* transcript isoform detected by snRNAseq of *Drosophila* testis tissue (SOM Figure 10). Western Blots of testes from flies homozygous for the *HA:cbc* allele probed with anti-HA showed expression of the expected ∼50 KDa isoform (Figure 6 B).

Immunoprecipitation with anti-HA antibodies from extracts of testes from flies homozygous for *HA:cbc* co-immunoprecipitated the >250 KDa band from the PCF11 isoforms, consistent with the Cbc and PCF11 proteins interacting in a complex (Figure 6 C). The 70 KDa isoform of PCF11 was not detected in the immunoprecipitate (entire gels of Western Blots and co-IP experiments are reported in SOM 11). Consistent with physical interaction between the Cbc and PCF11 proteins, accumulation of Cbc protein in spermatocytes was at least partially sensitive to the levels of PCF11 protein. Analysis by RNA-seq showed that when PCF11 expression was knocked down in late spermatogonia and spermatocytes by RNAi under control of *bamGal4*, *cbc* transcript was still expressed in *PCF11* knock down testes at a similar level compared to control testes (Figure 6 D). Likewise, in testes in which *cbc* had been knocked down by RNAi under control of *bamGal4*, transcripts from the *PCF11* locus were still expressed (Figure 6 E). However, the situation was different at the level of protein accumulation in spermatocytes. Simultaneous immunostaining of testis from *HA:cbc/+;bamGal4* males with anti-PCF11 and anti-HA antibodies showed both proteins present in spermatocyte nuclei from control males (Figure 6 F). However, for two different RNAi lines directed against *PCF11*, knock down of *PCF11* under control of *bamGal4* resulted in drastic reduction of signal from HA:Cbc (Figure 6 G’-H’, quantified in K). Similar reduction of Cbc signal in *UAS>PCF11_VDRC-RNAi;bamGal4* and *UAS>PCF11_NIG-RNAi;bamGal4* spermatocytes was observed in flies homozygous for the *HA:cbc* tagged allele (SOM 12). In contrast, knock down of *cbc* by RNAi under control of *bamGal4* did not cause loss of signal from anti-PCF11 (Figure 6 I’’-J’’, quantified in L).

### Forcing high expression of *cbc* and/or *PCF11* in spermatogonia induced cleavage at proximal PASs normally used in spermatocytes

Our data from the knock down experiments in spermatocytes suggest that the high levels of CFII components expressed in spermatocytes are required for cleavage of nascent transcripts of certain genes at an upstream PAS rather than at the distal PAS used in spermatogonia. Forced overexpression of CFII components in spermatogonia triggered a fraction of the nascent transcripts from certain genes to be cleaved at the upstream site normally used in spermatocytes.

Minigenes with the coding sequences of *cbc* with a V5 tag or *PCF11-RF* with an HA-FLAG tag, both under the control of two repeats of 5X UAS and the *hsp70* promoter, were introduced into the *Drosophila* genome. These transgenes allow spermatogonial specific overexpression of *cbc-V5* and/or *PCF11-RF-HA* by crossing males carrying the designed UAS constructs with females carrying a combination of *nanosGal4* (to drive in stem cells and early spermatogonia) and *bamGal4* (to drive in mid-late spermatogonia). Transgenes and Gal4 expressing lines were introduced into a *bam*^-/-^ genetic background, in which early stage spermatogonia over proliferate and fail to differentiate into spermatocytes and expression from the transgenes scored by immunofluorescence staining with anti-V5 and anti-HA (Figure 7 A-D). Anti-HA staining to visualize the overexpressed PCF11-RF-HA showed that the protein was highly nuclear in spermatogonia from flies overexpressing *PCF11-HA-RF* (Figure 7 B’’) and from flies overexpressing simultaneously *cbc-V5* and *PCF11-RF-HA* (Figure 7 D’’), recapitulating PCF11 nuclear localization in wild-type spermatogonia (Fig 1 J). Anti-V5 staining to visualize the overexpressed Cbc-V5 was instead both nuclear and cytoplasmic in spermatogonia from transgenic flies overexpressing *cbc-V5* (Figure 7 C’). However, in spermatogonia overexpressing both *PCF11-RF-HA* and *cbc-V5*, more of the anti-V5 signal was nuclear and less cytoplasmic (Figure 7 D’). Quantification of the fluorescent signal from the nucleus and cytoplasm showed that the ratio of anti-HA signal in the nucleus compared to the cytoplasm remained constant when *PCF11-RF-HA* was overexpressed alone or with *cbc-V5*. However, the ratio of nuclear to cytoplasmic anti-V5 signal increased when *cbc-V5* was overexpressed with *PCF11-RF-HA* compared to when *cbc-V5* was overexpressed alone (Figure 7 E). Immunoprecipitation of Cbc-V5 using anti-V5 antibody from extracts of *bam*^-/-^ testes in which *PCF11-RF-HA* and *cbc-V5* were both overexpressed in spermatogonia under control of *nanosGal4* and *bamGal4* brought down PCF11-RF-HA, indicating that the overexpressed tagged proteins interact, similar to their counterparts expressed in wild-type testes (Figure 7 F, entire gels in SOM Figure 13).

**Figure 7.**
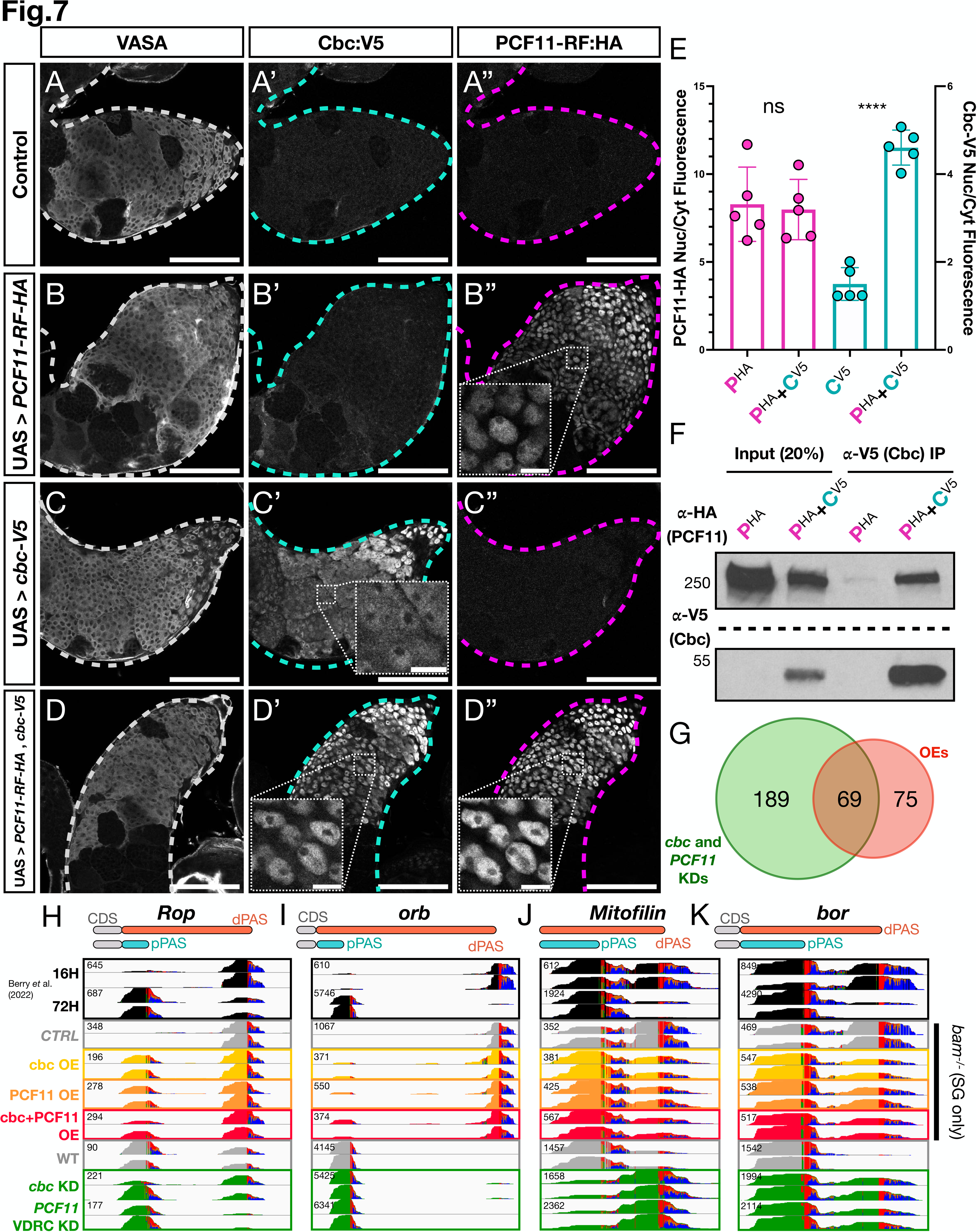
Overexpression of PCF11 and Cbc in spermatogonia shifted cleavage towards the more proximal site for some transcripts. (A-D) *bam*^-/-^ testes stained with anti-Vasa, anti-HA and anti-V5. (A-A’’) *bam* mutant control lacking overexpression constructs (*nosGal4/Y; +/CyO; bam*^86^*,bamGal4/bam*^1^), (B-B’’) *bam* mutant testes overexpressing *PCF11-RF-HA (nosGal4/Y; UAS>PCF11-RF-HA/+; bam*^86^*,bamGal4/bam*^1^), (C-C’’) *bam* mutant testes overexpressing *cbc-V5 (nosGal4/Y; UAS>cbc-V5/+; bam* ^86^*,bamGal4/bam*^1^), (D-D’’) *bam* mutant testes overexpressing both *PCF11-RF-HA* and *cbc-V5 (nosGal4/Y; UAS>PCF11-RF-HA, UAS>cbc-V5/+; bam*^86^*,bamGal4/bam*^1^). (E) Pink: quantification of nuclear/cytoplasmic ratio of PCF11-RF-HA in *bam*^-/-^ testes overexpressing *PCF11-RF-HA* alone or *PCF11-RF-HA* plus *cbc-V5*. Blue: quantification of nuclear/cytoplasmic ratio of cbc-V5 in *bam*^-/-^ testes overexpressing *cbc-V5* alone or *PCF11-RF-HA* plus *cbc-V5*. For each sample N=5 testes were quantified. Statistical analysis was performed using the Welch’s t-test. p-values are 0.8103 (ns) and <0.0001 (****). (F) Immunoprecipitation (IP) of *cbc-V5* from lysates of *bam*^-/-^ testes in which either *PCF11-RF-HA* plus *cbc-V5* or *PCF11-RF-HA* alone (control) were overexpressed, followed by PAGE and western blot probed with anti-HA and anti-V5 antibodies. Input = 20% of lysate before IP. (G) Venn diagram showing the number of genes that are affected in opposite APA directions by CFII knock downs (in green) or CFII overexpression (in red). The green circle represents all the genes that had a polyA index > 0.5 (lengthening) in either *PCF11* or *cbc* knock downs, while the red circle all the genes that had a polyA index < -0.5 (shortening) in either *PCF11-RF-HA* overexpression, or *cbc-V5* overexpression, or *PCF11-RF-HA* plus *cbc-V5* overexpression. (H-K) IGV plots showing 3’ end seq reads from libraries for overexpression and knock down samples for the (H) *Rop,* (I) *orb,* (J) *Mitofilin* and (K) *bor* loci. OE=overexpression, SG=spermatogonia. (CTRL) = *nosGal4/Y; +/+; bam*^86^*,bamGal4/bam*^1^, (*cbc* OE) *= nosGal4/Y; UAS>cbc-V5/+; bam*^86^*,bamGal4/bam*^1^*, (PCF11* OE) *= NosGal4/Y; UAS>PCF11-RF-HA/+; bam* ^86^*,bamGal4/bam*^1^*, (cbc + PCF11* OE) = *nosGal4/Y; UAS>PCF11-RF-HA, UAS>cbc-V5/+; bam*^86^*,bamGal4/bam*^1^.

Analysis of 3’seq libraries from *bam*^-/-^ testes in which *cbc-V5* alone, *PCF11-RF-HA* alone, or *cbc-V5* and *PCF11-RF-HA* together were overexpressed in spermatogonia compared with 3’seq libraries from *bam*^-/-^ testes in which neither was overexpressed revealed that of the ∼500 genes identified by Berry *et al*. (2022) as producing mRNAs with different 3’UTR length due to APA, ∼140 showed upregulation of 3’ end cleavage at the proximal PAS (see for example the genes *Rop, orb, Mitofilin* and *bor* (Figure 7 H-K). 47% of the genes that showed an increase in usage of the proximal PAS normally used in spermatocytes (polyA index < -0.5) in samples in which either *cbc-V5* or *PCF11-RF-HA* or both were overexpressed in *bam*^-/-^ spermatogonia also showed higher usage of the distal PAS in samples in which *cbc* or *PCF11* had been knocked down in spermatocytes (Figure 7 G). The increased processing at proximal PASs in *bam*^-/-^ testes in which CFII components were overexpressed only affected some of the nascent transcripts from a given locus and was never an APA switch as striking as the one observed when *cbc* or *PCF11* were knocked down in spermatocytes (SOM Figure 14 and Figures 3 and 4).

## Discussion

Recent results show that stage-specific Alternative Cleavage and Polyadenylation (APA) can regulate dramatic changes in the suite of proteins expressed as cells advance stepwise through a differentiation sequence (Berry *et al*. 2022). The developmental mechanisms that regulate cell type and stage-specific APA can thus be key contributors to cellular differentiation. Our analysis has revealed that cell-type-specific differential expression of components of Cleavage Factor complex II (CFII) of the 3’ end cleavage machinery regulates alternative polyadenylation in the *Drosophila* male germ line adult stem cell lineage. Results of cell-type-specific knock down experiments suggest that the developmentally specified increase in expression of CFII components PCF11 and Cbc is important for the shift to 3’ end cleavage at a more proximal site in spermatocytes compared to spermatogonia at over 200 loci during *Drosophila* spermatogenesis. Forcing overexpression of CFII components in spermatogonia resulted in increased 3’ end cleavage at the proximal site more typical of spermatocytes for a fraction of transcripts from some genes. Knock down of several other cleavage factors in spermatocytes had much milder effects on shifting the 3’ end cleavage from the proximal site to the more distal site normally utilized in spermatogonia, singling out CFII for a key role in the developmentally regulated 3’UTR shortening due to APA characteristic of wild-type *Drosophila* spermatocytes.

Cell-type-specific upregulation of *cbc* may be a key factor in developmental regulation of APA in differentiating male germ cells. While *PCF11* transcripts are broadly expressed in fly adult tissues, *cbc* seems to be particularly highly expressed mostly in spermatocytes. In addition, previous studies involving knock down of *PCF11* in cancer cells and zebrafish embryos showed that downregulating *PCF11* broadly induced 3’UTR lengthening. On the other hand, knocking down *CLP1* (the mammalian homolog of *cbc*) in mammalian cells did not show remarkable effects on APA (Li *et al*. 2015; Ogorodnikov *et al*. 2018; Kamieniarz-Gdula *et al*. 2019). However, it is possible that *cbc* was not highly expressed or did not play a key role in regulating APA in the cell types tested. In contrast, knock down of *cbc* in *Drosophila* spermatocytes strongly affected the APA profile, resulting in cleavage at the distal site typical of spermatogonia for almost half of the genes that would instead normally be cleaved at proximal sites in spermatocytes.

Several lines of evidence suggest that PCF11 may carry out functions independently of CFII. PCF11 has been found to bind the C-Terminal Domain (CTD) of RNA POLII *in vitro* and in yeast (Sadowski *et al*. 2003; Zhang and Gilmour 2006), where it acts as a termination factor (Guéguéniat *et al*. 2017; Schäfer *et al*. 2018; Kamieniarz-Gdula *et al*. 2019). ChiP-seq of POLII in control versus *PCF11* knock down human cells showed that POLII fails to terminate efficiently when PCF11 is reduced (Kamieniarz-Gdula *et al*. 2019). Accordingly, downregulating *PCF11* expression levels has been shown to induce 3’UTR lengthening in cancer cells and zebrafish embryos (Li *et al*. 2015; Ogorodnikov *et al*. 2018; Kamieniarz-Gdula *et al*. 2019). Whether skipping upstream cleavage sites might be linked with the dynamics of POLII termination, and what other functions PCF11 might carry out in association with RNA POLII still need to be elucidated. PCF11 has been shown to bind RNA *in vitro*, with a preference for G-rich regions. On the other hand, Cbc does not have a recognized RNA binding domain, and immunoprecipitation of Cbc *in vitro* failed to detect bound RNA (Schäfer *et al*. 2018). Our findings raise the possibility that Cbc might act as a developmentally regulated factor that, when upregulated in certain cell types, might bind to PCF11 to form CFII, utilize the ability of PCF11 to recognize RNA, and modulate PCF11 activity to enhance 3’ end cleavage at specific sites.

Notably, not all of the ∼500 genes found to undergo cell-type-specific 3’UTR shortening due to APA in *Drosophila* spermatocytes were affected by *PCF11* or *cbc* knock down, suggesting that other developmental mechanisms (such as changes in transcription rates and/or activity of other RNA binding proteins) might play a role in regulating stage-specific APA at some genes. In addition, forced overexpression of CFII components in spermatogonia increased 3’ end cleavage at the proximal PAS normally used in spermatocytes only for some genes, suggesting that additional factors (perhaps PABP2, for example) may help drive cleavage at the proximal PAS. It is also possible that CFII has spermatocyte specific components or interacting proteins that still need to be identified.

PCF11 has been found to be part of an autoregulatory feedback loop in cancer cells and zebrafish, where it auto-regulates its own protein levels by activating a premature PAS (Kamieniarz-Gdula *et al*. 2019). Our 3’ end sequencing data revealed that a PAS located very far 5’ in the *PCF11-RC* coding sequence becomes highly utilized in spermatocytes compared to spermatogonia. Interestingly, cleavage at this premature PAS was dependent on *PCF11* and *cbc* levels, as the upstream PAS was no longer highly utilized if *PCF11* or *cbc* were knocked down in spermatocytes (SOM Figure 15). This could mimic what has been observed in human cells and zebrafish embryos and suggests that the proposed PCF11 auto-regulatory loop might be dependent on both components of the CFII complex rather than on PCF11 alone.

Developmentally regulated switches in the site at which nascent mRNAs are terminated, such as those we have observed to be dependent on components of CFII, may serve as a mechanism to quicky change the menu of proteins expressed as cells progress through sequential steps in differentiation. By effecting the length and content of 3’UTRs, APA can change whether mRNAs are translated, blocked from translation, differentially localized, stabilized, or degraded. Since APA is a co-transcriptional process that does not require opening or closing of chromatin or re-initiation of transcription, it may provide a more rapid way to turn expression of some proteins on and other proteins off, changing cell state and canalizing cell fate decisions. As such, understanding how the alternative choices in 3’ end cut site are regulated is key for understanding developmental regulation of cell states.

## Materials and Methods

### Fly Strains, husbandry and design of UAS transgenes for overexpression

Flies were grown on standard molasses media and kept at 25 °C, 29 °C or room temperature as indicated. For RNAi induced knock downs, fly crosses (*bamGal4* driver females X RNAi line males) were kept at 25 °C for three days. Adults were then shifted to a new media bottle and kept at 25 °C while the eggs and early larvae were shifted to 29 °C to increase knock down efficiency. To further increase knock down penetrance, we used *Gal4* drivers that also expressed *UAS > dicer2* . For over expression in spermatogonia of *PCF11* and *cbc*, male flies expressing minigenes of *PCF11-RF*-HA-FLAG and/or *cbc*-V5 under the control of two 5XUAS sequences and a *hsp70* promoter were crossed with females bearing both the *nanos*Gal4 and *bam*Gal4 transgenes to induce expression in both early (*nanosGal4*) and late (*bamGal4*) spermatogonia. Crosses were kept three days at 25 °C. Then the adults were removed, and the eggs and larvae shifted to 29 °C.

Plasmid manipulation was performed following standard cloning strategies. In brief, for the plasmid to force expression of *PCF11-RF-HA*, the plasmid LD11480 from DGRC (Drosophila Genomics Resource Center) was used as a template, while for *cbc-V5* forced expression the plasmid UFO01678 from DGRC was used as a template. Plasmid injection and final line balancing were performed by BestGene. The construct to overexpress *PCF11-RF-HA* was inserted on an attP landing site on the left arm of the second chromosome, while the construct to overexpress *cbc-V5* was inserted on an attP landing site on the right arm of the second chromosome. Plasmid maps are reported in supplementary material. Fly lines used in this study: *W1118* (BL #3605), *bamGal4* (Chen and McKearin 2003), *nanosGal4-VP16* (Doren *et al*. 1998), *bam*^1^/TM6B (McKearin and Spradling 1990), *bam*^86^/TM6B (McKearin and Ohlstein 1995), *CPSF6:GFP* (VDRC #318105), *HA:Cbc* (N-terminal CRISPR tag, kindly gifted by the Wang lab, (Wu *et al*. 2021)), *PCF11*-RNAi_1 (VDRC #103710) *PCF11*_RNAi_2 (NIG #10228R4), *cbc*-RNAi_1 (BL #77426), *cbc*-RNAi_2 (VDRC #100686), *CPSF6*-RNAi_1 (BL #34804), *CPSF6*-RNAi_2 (VDRC #107147), *CstF64*-RNAi_1 (BL #65987), *CstF50*-RNAi_1 (BL #77377), *PABP2*-RNAi_1 (VDRC #106466), *PABP2*-RNAi_2 (VDRC #33499), *su(f)*-RNAi_1 (BL 55693), UAS > *PCF11-RF*-HA-FLAG (this study), UAS > *cbc*-V5 (this study).

### Immunofluorescence (IF) staining and signal quantification

Testes from 0-3 days old males were removed by dissecting in 1X PBS on a cyclops dissecting dish at room temperature over 30 minutes or less, then testes were immediately fixed in a 1.7 mL Eppendorf tube with 1 mL 5% formaldehyde in 1X PBS for 30 minutes at room temperature with rocking. If needed, fixed samples were stored in 1X PBS at 4 °C for up to one week. Testes were then permeabilized in 0.3% Triton-X-100 and 0.6% sodium deoxycholate in 1X PBS for 30 minutes at room temperature, rinsed twice in 1X PBS + 0.05% Tween-20 (PBT), blocked for 30 minutes in 1X Western Blocking Reagent (WBR, Roche #11921673001) in PBT at room temperature, then incubated with primary antibody mixed in 1X WBR+PBT overnight at 4 °C. After primary antibody incubation, testes were rinsed twice in PBT, blocked again for 30 minutes in 1X WBR+PBT at room temperature, then incubated with secondary antibody mixed in 1X WBR+PBT for 2 hours at room temperature in a dark container. Testes were then rinsed twice in PBT and mounted on a glass slide in 10 uL of ProLong diamond anti-fade with DAPI (ThermoFisher, CAT #P36962). Primary antibodies used: guinea pig anti-CstF50 (1:300, kindly gifted by John T. Lis (Ni *et al*. 2008)), rabbit anti-PABP2 (1:500, kindly gifted by Martine Simonelig (Benoit *et al*. 1999)), goat anti-Vasa (1:200, Santa Cruz Biotechnology sc-26877, discontinued), rat anti-Vasa (1:10 for supernatant or 1:100 for the concentrated version, both from DSHB, antibody ID 760351) rabbit anti-CstF64 (1:400, kindly gifted by Zbigniew Dominski (Skrajna *et al*. 2018)), rabbit anti-PCF11 (1:200, kindly gifted by David Scott Gilmour (Zhang and Gilmour 2006)), rabbit anti-HA (1:500, Cell Signalling Technologies, #3724), mouse anti-HA (1:200, Biolegends #901501), chicken anti-GFP (1:10.000, Abcam #13970) and mouse anti-V5 (1:200, Invitrogen #46-0705). Secondary antibodies used: donkey or goat anti-rabbit, anti-mouse, anti-guineapig, anti-chicken, anti-rat and anti-goat, all used at 1:500 and conjugated with either Alexa 488, Alexa 568, Alexa 594 or Alexa 647 depending on the combination of antibodies used in each experiment (all from the ThermoFisher catalog). Imaging of IF experiments was performed on a Leica SP8 Confocal microscope. Protein levels in Figures 1 and 6 were quantified in Fiji by manually delineating the cell nuclei and applying the formula: *Integrated_Density – (Area X Background_Mean)*. The Nuclear-Cytoplasmic ratio of Cbc-V5 and PCF11-RF-HA proteins in Figure 7 were quantified in Fiji by measuring the mean intensity of protein signal in 10 different uniform pixels areas per image in the central region of the sample and applying the formula: *(Mean_protein – Mean_background)* _Nucleus_ */ (Mean_protein – Mean_background)*_Cytoplasm_.

### HCR^TM^-RNA-FISH

Testes were dissected from 0-3 day old males in 1X PBS on a cyclops dissecting dish at room temperature over a maximum of 30 minutes, then fixed and permeabilized as done for immunofluorescence. The protocol for “generic sample in solution” from the Molecular Instruments website (https://www.molecularinstruments.com/hcr-rnafish-protocols) was followed. Hybridization buffer, wash buffer, amplification buffer and hairpins were all from Molecular Instruments (HCR™ RNA-FISH Bundle). After permeabilization as described for immunofluorescence above, samples were incubated with 200 uL of probe hybridization buffer for 30 minutes at 37 °C. Probe mixes were prepared by mixing 3.2 uL of 1 uM gene-specific-probes mix (one probe mix per gene or transcript) in 200 uL of probe hybridization buffer. Hybridization buffer was removed, probe mix added, and the samples were incubated overnight in probe mix at 37 °C in a water bath. The following day samples were washed 4 times for 15 minutes each in 200 uL of probe wash buffer at 37 °C and then washed 3 times 5 minutes each in 5X SSCT [5X sodium chloride sodium citrate (SSC, Invitrogen #AM9770) and 0.1% Tween-20 in H_2_O] at room temperature. Samples were then incubated in 500 uL of amplification buffer for 30 minutes at room temperature. In the meantime hairpin mix was prepared: 30 pmol per sample (5 uL) of harpin h1 and hairpin h2 (each fluorophore has its own harpin combination, in our case since we always imaged 2 transcripts at the same time, we used 2 fluorophores and therefore 4 hairpins) were heated at 95 °C for 90 seconds in a thermocycler in the dark and cooled down at room temperature in a drawer for 30 minutes (each hairpin was heated up and cooled in a separate tube). To prepare hairpin mixes 5 uL of each harpin were added to 250 uL of amplification buffer. The samples were then incubated in harpin mix at room temperature in the dark overnight in a drawer with no rocking. The following day samples were washed with 500 uL 5X SSCT 2 times for 5 minutes each, then 2 times 30 minutes each and finally 1 time 5 minutes, always in the dark and at room temperature. Samples were then mounted on a glass slide in 10 uL of ProLong diamond anti-fade with DAPI (ThermoFisher, CAT #P36962). Probes were designed as in Bedbrook *et al*. 2023 and ordered from Integrated DNA technologies as Oligo Pools (oPools^TM^ 50 pmol/oligo). Harpins H1 and H2 were ordered conjugated with either 488 or 546 fluorophores from Molecular Instruments. Probe sequences are listed in supplementary material. Imaging was performed on a Leica SP8 confocal microscope. For each testis, 5 areas of spermatocyte cytoplasm from different testis regions were quantified on Fiji from both probe channels, the background was subtracted and then the ratio between the probe recognizing the 3’UTR extensions and the probe recognizing the protein coding region was calculated. The following formula was used: *(Area_Mean_p2 – Back_Mean_p2) / (Area_Mean_p1 – Back_Mean_p1)* where p2 is the probe set recognizing the 3’UTR extension while p1 is the probe set recognizing the protein coding region of the transcript. Area’s size varied between pictures but was constant between the 5 measurements from the same image.

### 3’ end sequencing and analysis

Testes were dissected from 0-3 day old males in S2 media in batches of 20-80 flies in a cyclops dissecting dish at room temperature over a maximum of 1 hour per batch. Testes in the batch were then together transferred to a 1.7 mL Eppendorf tube containing 700 uL 1X PBS. The 1X PBS was then immediately removed and the testes snap-frozen in liquid nitrogen and stored at -80 °C. RNA was extracted from ∼150 testes pairs using the RNAeasy Mini kit (Qiagen, 74104). Frozen tissue was dissociated using a 1mL syringe with a 27G needle aspirating up and down ∼10 times in RNAeasy Mini kit lysis buffer supplied with 1:100 β-mercaptoethanol as suggested by the kit manual. The quality of extracted RNA was assessed via bioanalyzer, and the concentration measured by Nanodrop and Qubit RNA HS Assay Kit (Life Technologies, Q32852). 3’ end RNAseq libraries were prepared using the Quant Seq (FWD) kit from Lexogen, using 500 ng of total extracted RNA per replicate. The quality of DNA libraries was checked via bioanalyzer, and the concentration measured with Qubit dsDNA HS Assay Kit (Life Technologies, Q32854). Sequencing was performed by the Stanford Genomics facility on an Illumina NextSeq 500 platform as in Berry *et al*. (2022). Each biological sample was performed in duplicate. Reads were filtered to retain only reads with 10 consecutive or more ‘A’. Reads were then trimmed to remove adaptor sequences and aligned to the *Drosophila* dm6 genome using bowtie 0.12.8 with a value of e=5000 to allow for the expected mismatch in the PolyA tail sequence, as in Berry *et al*. (2022). Quantification of PAS usage for the ∼500 genes identified in Berry *et al*. 2022 was performed using PolyA miner (Yalamanchili *et al*. 2020), using as input a bed file containing the proximal and distal PAS of the ∼500 genes identified by Berry *et al*. (2022) as cleaved at a proximal PAS in spermatocytes and a distal PAS in spermatogonia. To consider various spermatocyte stages we generated a master bed file by merging the proximal and distal cleavage sites from 3 different lists: *bam*^-/-^ (enriched for spermatogonia) vs. 32H (very early spermatocytes), *bam*^-/-^ vs. 48H (polar spermatocytes) and *bam*^-/-^ vs 72H (mid-stage maturing spermatocytes). 32H, 48H and 72H indicate hours after heat shock, which causes a pulse of bam expression in *bam*^-/-^ flies, inducing differentiation, as explained in Kim *et al*. (2017). As in these lists the coordinates of the cleavage sites did not match perfectly, some genes had more than 2 cleavage sites considered. However, PolyA miner was devleoped to consider genes with multiple polyA sites.

### *De novo* PAS calling and APA analysis

For analysis of changes in PAS usage genome wide between wild-type and knock down samples, we adapted the analysis pipeline from Zeng *et al*. 2024. The sequenced 3’ end-seq libraries were adapter-trimmed, and quality filtered using bbduk. The filtered reads were mapped to the dm6 *Drosophila* genome using STAR, then extracted for unique-mapping reads using Samtools. If a read was mapped to a region that was immediately upstream of six consecutive genomically encoded As or of a 10-bp region with at least 60% As, it was considered mis-primed and removed. The resulting filtered reads were used for *de novo* PAS calling using LAPA^61^ with the default setting, except requiring a replication rate cutoff at 0.95. For each gene, lengthening or shortening was quantified by considering the ratio between the 2 highest changing PAS: the PAS that is more upregulated in the test sample versus control sample and the PAS that is more downregulated in the test sample versus control sample. For this analysis we considered only genes for which both PASs located in the 3’UTR.

### RNAseq and analysis

The same total RNA extractions utilized for 3’seq was also used for preparation of short read RNAseq libraries. 1 ug of total extracted RNA was used to prepare libraries following the protocol from the NEBNext Ultra II Direction RNA Library Prep Kit for Illumina, following the section for PolyA selected RNA library preparation. Each biological sample was performed in duplicate. The quality and concentration of the libraries were measured via Bioanalyzer and Qubit, as for 3’seq libraries. Sequencing was performed by Novagene on a NOVAseq, PE 150 Illumina Platform. Raw reads were trimmed to remove the Illumina adapters using trim_galore, then aligned to the *Drosophila* dm6 genome using STAR. Differential expression analysis was performed using the package DEseq2.

### Fly Cell Atlas single nuclear RNAseq analysis

Using the Fly Cell Atlas “Testis” polydT-primed 10x single-nuclear RNA sequencing dataset (Li *et al*. 2023), for each of the 111 Leiden 6.0 clusters (as in Berry *et al*. 2022) we aggregated all reads from nuclei in that cluster. For each cluster, reads that uniquely mapped to one of the isoform specific *PCF11* 3’UTRs were counted and normalized by the length of that 3’UTR as well as the total number of reads derived from that cluster. This produced an isoform-specific rpkm value for each Leiden 6.0 cluster that was then log-normalized and plotted using Seurat. Gene expression and cell type annotation plots across all the adult *Drosophila* tissues analyzed in the Fly Cell Atlas project were generated in Seurat using published code modified to visualize relevant genes and cell identities of interest (Raz *et al*. 2023).

### Western Blots

Testes from 20-50 (depending on the genotype and the protein needed to be detected) 0-3 days old males were quickly dissected into 1X PBS in a cyclops dissecting dish at room temperature, transferred to a 1.7 mL Eppendorf tube containing 700 uL 1X PBS. The 1X PBS was then removed and the tube snap frozen in liquid nitrogen and stored at -80 °C until needed. Frozen tissue was lysed in 40 uL of cold Lysis Buffer (either RIPA buffer [50 mM TRIS pH 7.5, 250 mM NaCl, 0.1% SDS, 0.5% Sodium deoxycholate to which was added one tablet per 50 uL of protease inhibitor cocktail (cOmplete ULTRA Tablets, Mini, EDTA-free, EASY pack, Roche, REF 05 892 791 001)] or JUME Buffer [10 mM MOPS, 10 mM EDTA, 8M UREA, 1% SDS]). Tissue was dissociated using a 1 mL syringe with a 27G needle aspirating up and down ∼10 times, then incubated rocking at 4°C for 20 minutes. Following incubation, lysate was centrifuged at 15000 rpm for 5 minutes at 4°C. The supernatant was transferred to a new tube, mixed with 4X Laemli Buffer with β-mercaptoethanol, boiled for 7 minutes at 85 °C, then run on a Mini-PROTEAN TGX Precast Protein Gel (4–15% Mini-PROTEAN® TGX™ Precast Protein Gels, 10-well, 50 µl #4561084). The separated proteins were transferred from the gel to a PVDF membrane (BIO-RAD, Cat.# 1620177) overnight at 4°C. The following day, the membrane was blocked overnight at 4°C in 5% nonfat milk powder dissolved in PBST (1X PBS with 0.1% Tween-20). After blocking, the membrane was incubated for 1 hour at room temperature with primary antibody mix in 2% nonfat milk powder dissolved in PBST, washed 3 times 20 minutes each with PBST at room temperature, incubated with secondary antibody mix for one hour at room temperature in 2% nonfat milk powder dissolved in PBST and washed with PBST again 3 times 20 minutes each at room temperature. Bound antibodies were then revealed using Revvity Health Sciences Inc Western Lightning Plus-ECL, Enhanced Chemiluminescence Substrate (PerkinElmer #NEL104001EA). Primary antibodies used for Western blots: Rabbit anti-HA (1:5000 or 1:1000, Cell Signaling Technologies, #3724), Rabbit anti-PCF11 (1:10.000, kindly gifted from David Scott Gilmour (Zhang and Gilmour 2006)), mouse anti-V5 (1:1000, Invitrogen #46-0705). Secondary antibodies used for Western blots: Peroxidase AffiniPure™ Goat Anti-Rabbit IgG (H+L) (1:10.000, Jackson ImmunoResearch AB_2313567), Peroxidase AffiniPure™ Goat Anti-Mouse IgG (H+L) (1:10.000, Jackson ImmunoResearch AB_10015289).

### Immunoprecipitation (IP)

For Immunoprecipitation (IP) and co-IP experiments, testes were dissected into 1X PBS in a cyclops dissecting dish at room temperature in batches of 50 flies for a maximum of 30 minutes per batch, transferred into a 1.7 mL Eppendorf tube with 700 uL 1X PBS, the 1X PBS was removed and the tube snap-frozen in liquid nitrogen and stored at -80 °C. For one sample ∼100 pairs of testes were used. The day before the actual IP, 2 uL per sample of rabbit anti-HA antibodies (Cell Signaling Technologies, #3724) or 2uL of mouse anti-V5 antibodies (Invitrogen #46-0705), were crosslinked to magnetic beads (Dynabeads^TM^ Protein A Invitrogen #10002D for antibodies raised in rabbit, and Dynabeads^TM^ Pan Mouse IgG Invitrogen #11041 for antibodies raised in mouse) as described in Baker *et al*. (2023). Throughout this procedure, each of the following washes of the magnetic beads is done by gently resuspending beads in solution, placing the 1.7 uL containing beads Eppendorf tubes on a magnetic rack and removing the liquid. The day of the experiment, beads crosslinked with antibody were rinsed 1 time in 200 mM glycine pH 2.5 and washed once for 5 minutes in 200 mM glycine pH 2.5 to remove non-crosslinked antibodies, then blocked for 1 hour in 10% BSA in 50 mM Tris pH 8, and re-suspended in 20 uL of RIPA buffer and stored at 4 °C while the testis lysate was prepared. Frozen testis tissue was dissociated in RIPA buffer using a 1mL syringe and a 27G needle as in Western Blots experiments. The lysate was rocked at 4°C for 20 minutes, then centrifuged 5 minutes at max speed in a 1.7 mL Eppendorf tube at 4°C to separate lysate from tissue debris. The supernatant was transferred to a new 1.7 mL Eppendorf tube and incubated 45 minutes at 4 °C with beads with no antibody. Tubes containing lysate and beads were placed on a magnetic rack and 40 uL of precleared lysate was saved as input sample. Volume was brought back to 220 uL by adding 40 uL of cold RIPA buffer, pipetted into the tubes containing beads crosslinked to antibody and left incubating for 4 hours at 4°C with rocking. Since we noticed that HA:Cbc in testis lysates tended to form aggregates (see SOM Figure 11), we protein-precipitated (see protocol below) the input sample and resuspended the formed protein-pellet in JUME Buffer. 4X Laemli Sample Buffer (BIO-RAD #1610747) mixed with β-mercaptoethanol was added to the input sample to a final 1X concentration, incubated 7 minutes at 85 °C and frozen. After 4 hours rocking at 4°C, IP samples were placed on a magnet to separate beads from lysate and 40 uL of the supernatant were saved as supernatant sample, mixed with 4X Laemli Sample Buffer with β-mercaptoethanol incubated 7 minutes at 85°C and frozen. The rest of the supernatant was discarded. The beads with attached protein were immediately resuspended in RIPA buffer and washed with RIPA buffer 2 times for 5 minutes rocking at room temperature. Tubes were placed on a magnet to separate beads from RIPA buffer, then RIPA buffer was removed, and the beads were resuspended in 40 uL Elution Buffer (1X cOmplete ULTRA Tablets, 1% SDS, 10 mM EDTA, 50 mM Tris pH 8), vortexed, and incubated for 30 minutes at 70°C with frequent (every 5 minutes) vortexing or shaking. Tubes were then centrifuged at max speed for 10 seconds at room temperature, then placed on the magnet to separate beads from the eluate. 40 uL of eluate were moved to another 1.7 mL tube (IP-sample) to which was added 4X Laemli Sample Buffer with β-mercaptoethanol, then the sample was incubated 7 minutes at 85 °C and frozen. The beads were then mixed with 40 uL of elution buffer, mixed with 4X Laemli Sample Buffer with β-mercaptoethanol, incubated 7 minutes at 85 °C and frozen. Samples were then run on a Mini-PROTEAN TGX Precast Protein Gel and the protocol for Western Blots above followed. If the same membrane needed to be re-stained with another antibody, the membrane was stripped by incubation at 55 °C for 30 minutes in 62.5 mM Tris pH 6.8, 2% SDS and 100 mM β-mercaptoethanol, washed 3 times for 10 minutes each at room temperature in PBST and either stored in 1X PBS or blocked again before being re-stained as above. Alternatively, for a milder stripping, the membrane was incubated overnight at 4C with 5% nonfat milk in PBST + 0.1 mM sodium azide. The membrane was blocked again, then re-stained as above.

### Protein precipitation

To precipitate proteins, 4 volumes of ice-cold methanol were added to 1 volume of testis lysate, vortexed, and 1 volume of chloroform added. The sample was vortexed again, mixed with 3 volumes of H_2_O, vortexed and centrifuged in 1.7 mL Eppendorf tubes at 4 °C max speed for 5 minutes. After centrifugation, 2 liquid layers formed, with proteins at the interface. The upper liquid layer was removed, and 3 volumes of ice-cold methanol added to the sample. After a quick vortex, the sample was centrifuged at 4 °C max speed for 10 minutes to pellet the proteins. After centrifugation all liquid was removed, and the white protein pellet was air-dried and resuspended in 40 uL of JUME buffer.

## SOM Figure Legend

**SOM 1. Comparison of anti-HA staining between homozygous *HA:cbc/HA:cbc* and heterozygous *HA:cbc/CyO* testes.** (A) Apical testis from a male homozygous for *HA:cbc* and (B) a male *HA:cbc/CyO* heterozygous stained with anti-HA (Green) and DAPI (Blue). (A’-B’) show the HA:Cbc channel alone in grey corresponding to A and B. Scalebars are 100 uM. Green line marks spermatogonia region.

**SOM 2.** Heatmap related to Figure 4 showing more of the >500 genes from Berry et al. (2022) compared to the heatmap reported in Figure 4 C. Now are reported also genes that either have a PolyA index equal to zero in all the knock downs, a statistically non-significant PolyA index in all the knock downs, or a negative PolyA index in all the knock downs (that were omitted from the heatmap in Figure 4 C).

**SOM 3.** Volcano plots showing gene expression levels from RNAseq data from (A) *cbc-RNAi-BL 77426;bamGal4* and (B) *PCF11-RNAi-VDRC-103710;bamGal4* knock down testes versus *bamGal4* (control). In black are represented the genes that switch PAS between spermatogonia and spermatocytes from Berry *et al*. 2022 and that show a PolyA index either smaller than -0.5 or larger than 0.5 in the 3’seq libraries from the corresponding knock down.

**SOM 4. Verification of *PABP2* knock down efficiency** . (A, B) Apical tips of testes stained with anti-PABP2 antibodies and DAPI. (A) *bamGal4* (control) testes and (B) *PABP2-RNAi-VDRC-106466;bamGal4* testes. Arrow heads indicate somatic cyst cell nuclei, and a green line highlights the testis region with spermatogonia. (C-E) Phase contrast images of whole testes from: (C) Control: *bamGal4* testes, (D) *PABP2-RNAi-VDRC-106466;bamGal4* testes and (E) *PABP2-RNAi-VDRC-33499;bamGal4*. Scalebars are 50 uM (A, B) and 200 uM (C, D).

**SOM 5. Verification of knock down efficiency of CstF components** . Apical tips of testes from: (A) *bamGal4* (control) and (B) *CstF50-RNAi-BL-77377;bamGal4* males stained with anti-CstF50. (C) Phase contrast images of whole testes from *CstF50-RNAi-BL-77377;bamGal4* male. Apical tip of testes from (D) *bamGal4* (control) and (E) *CstF64-RNAi-BL-65987;bamGal4* testes stained with anti-CstF64 antibodies. (F) Phase contrast images of whole *CstF64-RNAi-BL-65987;bamGal4* testis. Arrow heads indicate somatic cyst cell nuclei, and a green line highlights the spermatogonia region. (G) Phase contrast microscopy of *su(f)-RNAi-BL-65693;bamGal4* testis. (H) Phase contrast images of whole *bamGal4* testis. Scalebars are 50 uM (A, B, D, E) and 200 uM (C, F, G, H).

**SOM 6. Knock down of *su(f)* or *CstF64* in spermatocytes alters CstF50.** (A, B) *bamGal4* (control) testis stained for (A) CstF64, and (B) CstF50. (C, D) *su(f-RNAi-BL-65693;bamGal4* testis and (E, F) *CstF64-RNAi-BL-65987;bamGal4* testis stained for (C, E) CstF64, and (D, F) CstF50. Scalebars are 100 uM.

**SOM 7. Verification of *CPSF6* knock down efficiency** . Phase contrast images of whole testes from (A-A’’’’) *CPSF6-RNAi-BL-34804;bamGal4* males, (B) a *CPSF6-RNAi-VDRC-107147;bamGal4* male and (C) a *bamGal4* male. Scalebars are 200 uM.

**SOM 8-9**. Changes in 3’ end cut site usage in knock down testes vs Control (*bamGal4*) testes on *de novo* mapped PAS sites on annotated 3’UTRs.

**SOM 10.** Umap from Fly Cell Atlas single nuclear RNA-seq (Raz *et al*. 2023; Li *et al*. 2023) showing relative expression of *PCF11-RC* in the testis and seminal vesicle sample.

**SOM 11.** Entire gels for Western Blots and co-IP from Figure 6.

**SOM 12. Cbc:HA is downregulated (but still present) upon *PCF11* knock down.** Apical tips of testes from (A, A’) *HA:cbc/HA:cbc;bamGal4* (B, B’) *HA:cbc,PCF11-RNAi-VDRC-103710/HA:cbc;bamGal4* (C, C’) *HA:cbc,PCF11-RNAi-NIG-10228R-4/HA:cbc;bamGal4* stained with antibodies against HA and PCF11. (D, E) Quantification of fluorescent signal from (D) PCF11 and (E) HA antibodies respectively in spermatogonia and spermatocytes as quantified in Figure 6. P-values are: (D) (*) = 0.0398 and (E) (*) = 0.0343 in *bamGal4* and (*) = 0.0438 in *VDRC 103710.* Scalebars are 100 uM.

**SOM 13.** Entire gels from co-IP in Figure 7

**SOM 14.** Heatmap reporting the PolyA index for genes from Berry et al. (2022) in 3’seq libraries from *bam*^-/-^ testes (filled with spermatogonia) overexpressing either *cbc-V5* alone, or *PCF11-RF-HA* alone, or botch *cbc-V5* and *PCF11-RF-HA*. Blue: proximal PAS more regulated in a sample overexpressing either *cbc-V5*, or *PCF11-RF-HA*, or both compared to control *bam*^-/-^ testes means that the percentage of transcripts cleaved at the proximal site typical of spermatocytes was higher ion the overexpression sample compared to *bam*^-/-^ testes in which neither *PCF11-RF-HA* nor *cbc-V5* was overexpressed.

**SOM 15.** IGV plots of 3’seq reads of *PCF11* locus. Gray arrow marks a PAS very 5’ in the locus that is utilized in samples enriched for spermatocytes but not much in samples enriched for spermatogonia. Cleavage at this upstream PAS is strongly downregulated in testes where either *PCF11* or *cbc* were knocked down in spermatocytes. SG = spermatogonia, SC = spermatocytes.

## Supporting information

HCR-probes-list

## Acknowledgements

We thank all the members of the Fuller lab for suggestions and comments on this manuscript and Dr. Nicole Martinez and Dr. Lauren Goins for their valuable feedback on the manuscript and suggested experiments. We thank Dr. Catherine Baker for suggestions on Immunoprecipitations and for proofreading the manuscript, and Benjamin Bolival for helping in maintaining fly crosses. We thank Dr. Jingxun Chen from the Brunet lab at Stanford for her help in designing HCR-FISH probes and for sharing some of the reagents, the Wang lab for the *HA:cbc* CRISPR tag flies, the Lis lab for anti-CstF50, the Simonelig lab for anti-PABP2, the Dominski lab for anti-CstF64 and the Gilmour lab for anti-PCF11 antibodies. We thank Daniel Mokhtari for his help parsing the Fly Cell Atlas dataset, and Dr. Hari Krishna Yalamanchili for his help in running PolyA miner. We thank the Cell Science Imaging facility at Stanford, supported, in part, by Award Number 1S10OD010580-01A1 from the National Center for Research Resources (NCRR) (The contents of this manuscript are solely the responsibility of the authors and do not necessarily represent the official views of the NCRR or the National Institutes of Health), particularly the CSIF facility manager Kitty Lee. We thank the Stanford Fly Media Center as well and all the staff working in the Developmental Biology department. L.G. was supported by an AICF (American Italian Cancer Foundation) postdoctoral fellowship (2021-2023). This research was supported by National Institutes of Health grant R35 GM136433) and funds from the Katherine D. McCormick and Stanley McCormick Memorial Professorship and the Reed-Hodgson Professorship in Human Biology to M.T.F. This manuscript is dedicated to Cuchuflí.

## Author Contributions

LG and MTF conceived the project and wrote the manuscript. LG designed the experimental work and performed or supervised all experimental and computational procedures. NRM contributed to experimental design and bioinformatic analysis. FMP contributed to experimental design and performed molecular cloning. IN performed immunofluorescence experiments. SS performed bioinformatic analysis of the Fly Cell Atlas data. YZ performed bioinformatic analysis to *de novo* map PASs. Funding was acquired by LG and MTF.

