## Supplementary Figures for "A Developmental Mechanism to Regulate Alternative Polyadenylation in an Adult Stem Cell Lineage"

**SOM Figure 1**

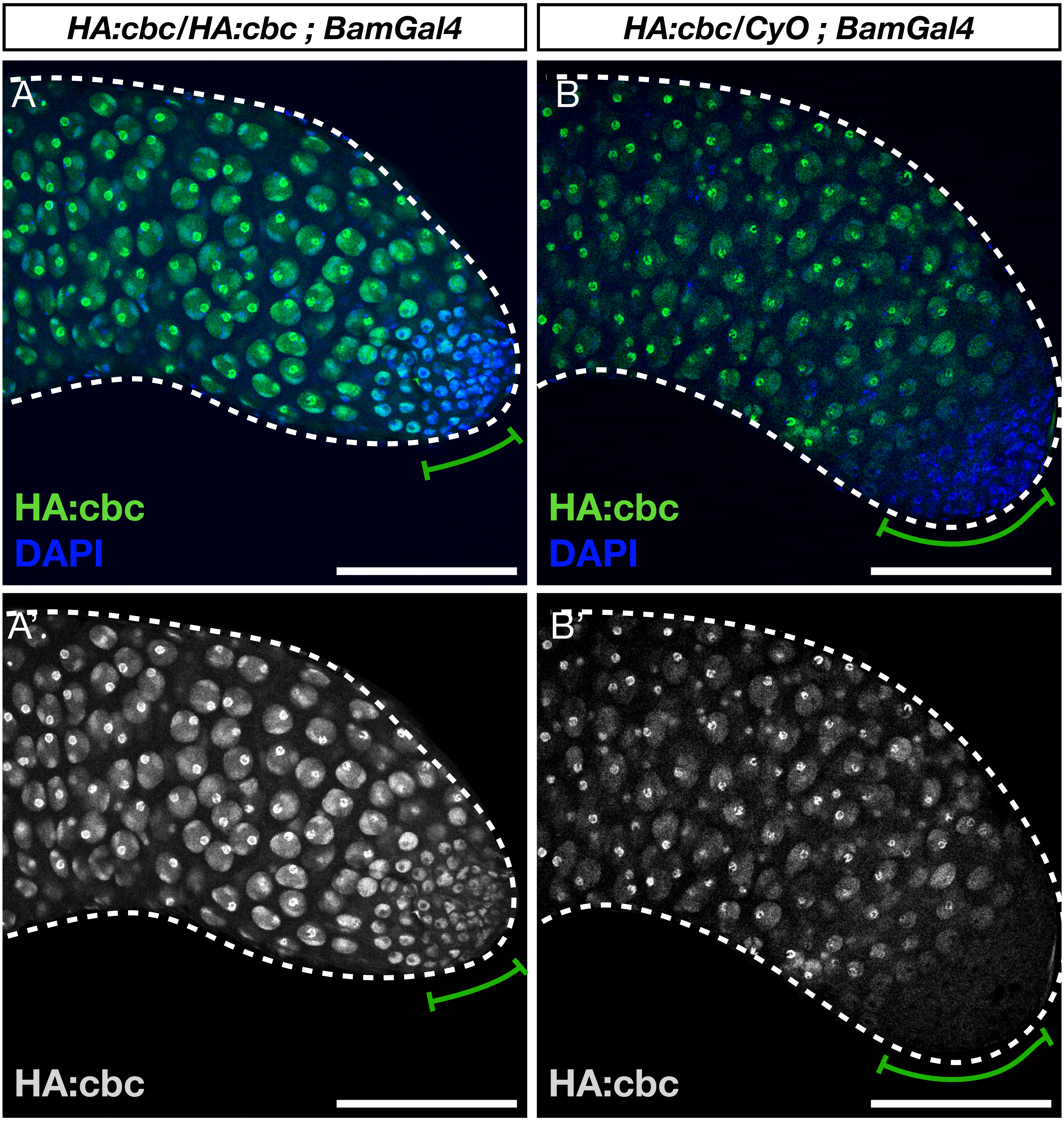
HA:cbc/HA:cbc ; BamGal4
HA:cbc/CyO ; BamGal4
A
HA:cbc
DAPI
B
HA:cbc
DAPI
A'
HA:cbc
B'
HA:cbc

### SOM Figure 2

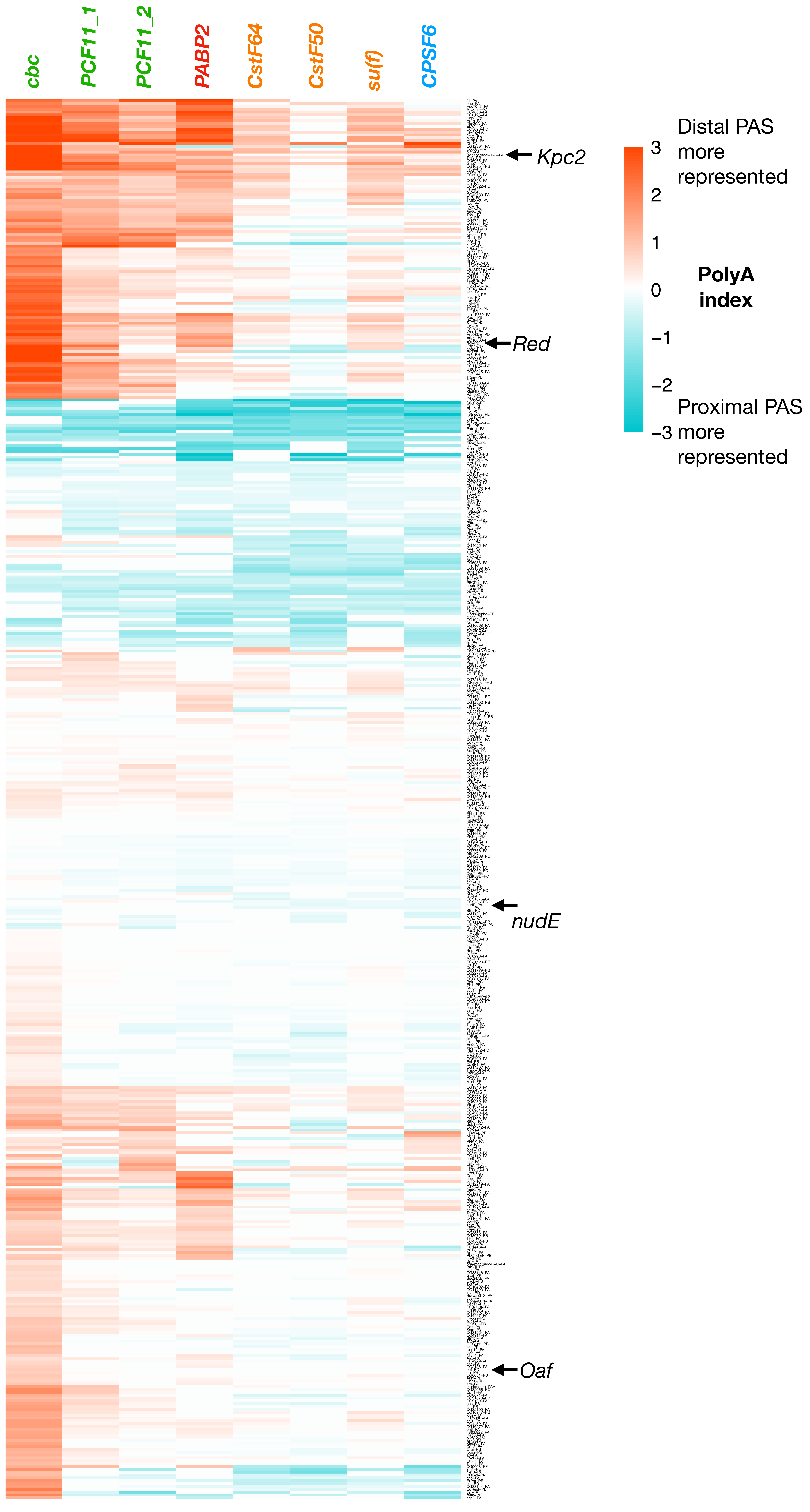
cbc
PCF11_1
PCF11_2
PABP2
CstF64
CstF50
su(f)
CPSF6
Kpc2
Red
nudE
Oaf
Distal PAS more represented
3
2
1
0
−1
−2
−3
PolyA index
Proximal PAS more represented

### SOM Figure 3

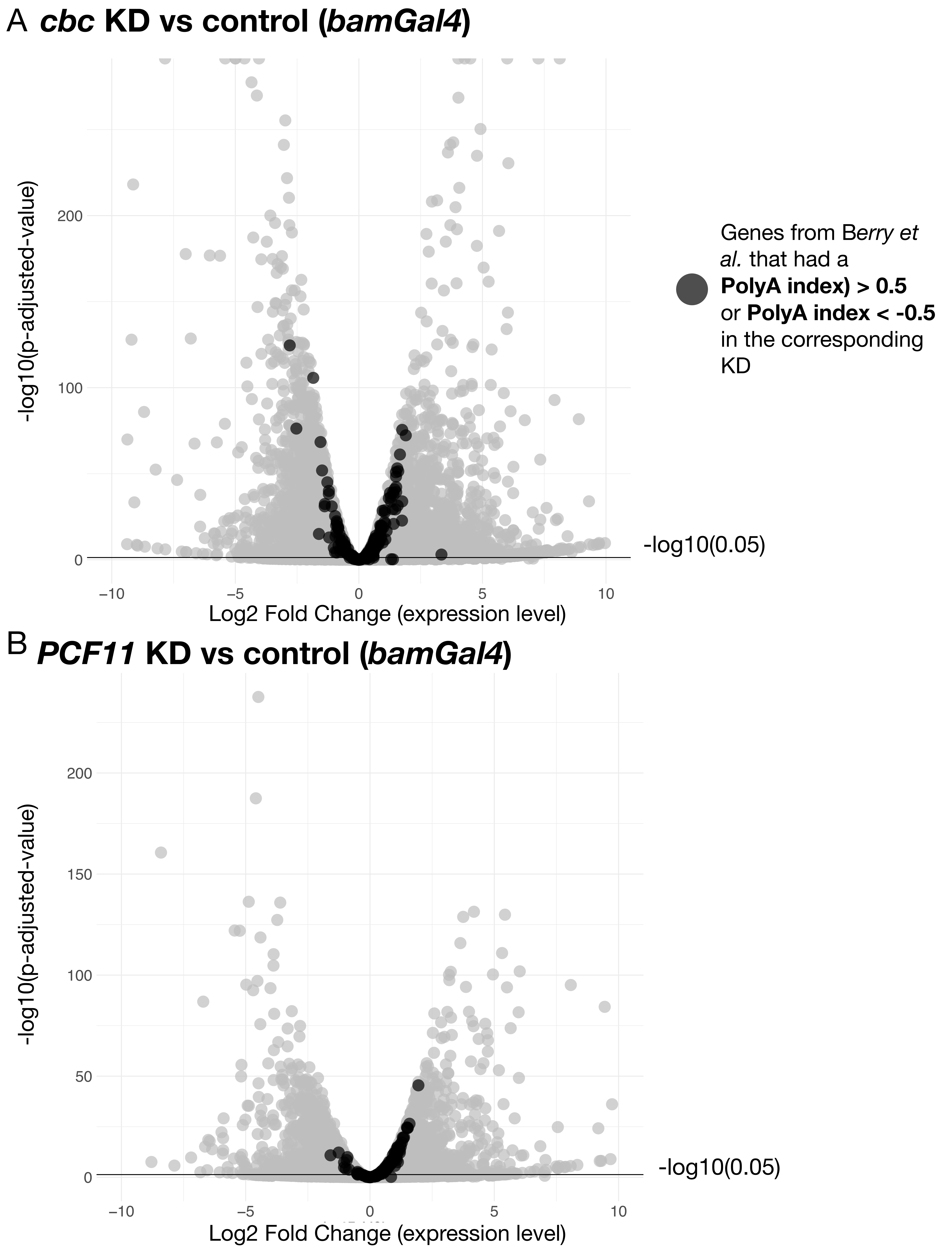
A cbc KD vs control (bamGal4)
-log10(p-adjusted-value)
200
100
0
-log10(0.05)
−10
−5
0
5
10
Log2 Fold Change (expression level)
Genes from Berry et al. that had a PolyA index) > 0.5 or PolyA index < -0.5 in the corresponding KD
B PCF11 KD vs control (bamGal4)
-log10(p-adjusted-value)
200
150
100
50
0
-log10(0.05)
−10
−5
0
5
10
Log2 Fold Change (expression level)

### SOM Figure 4

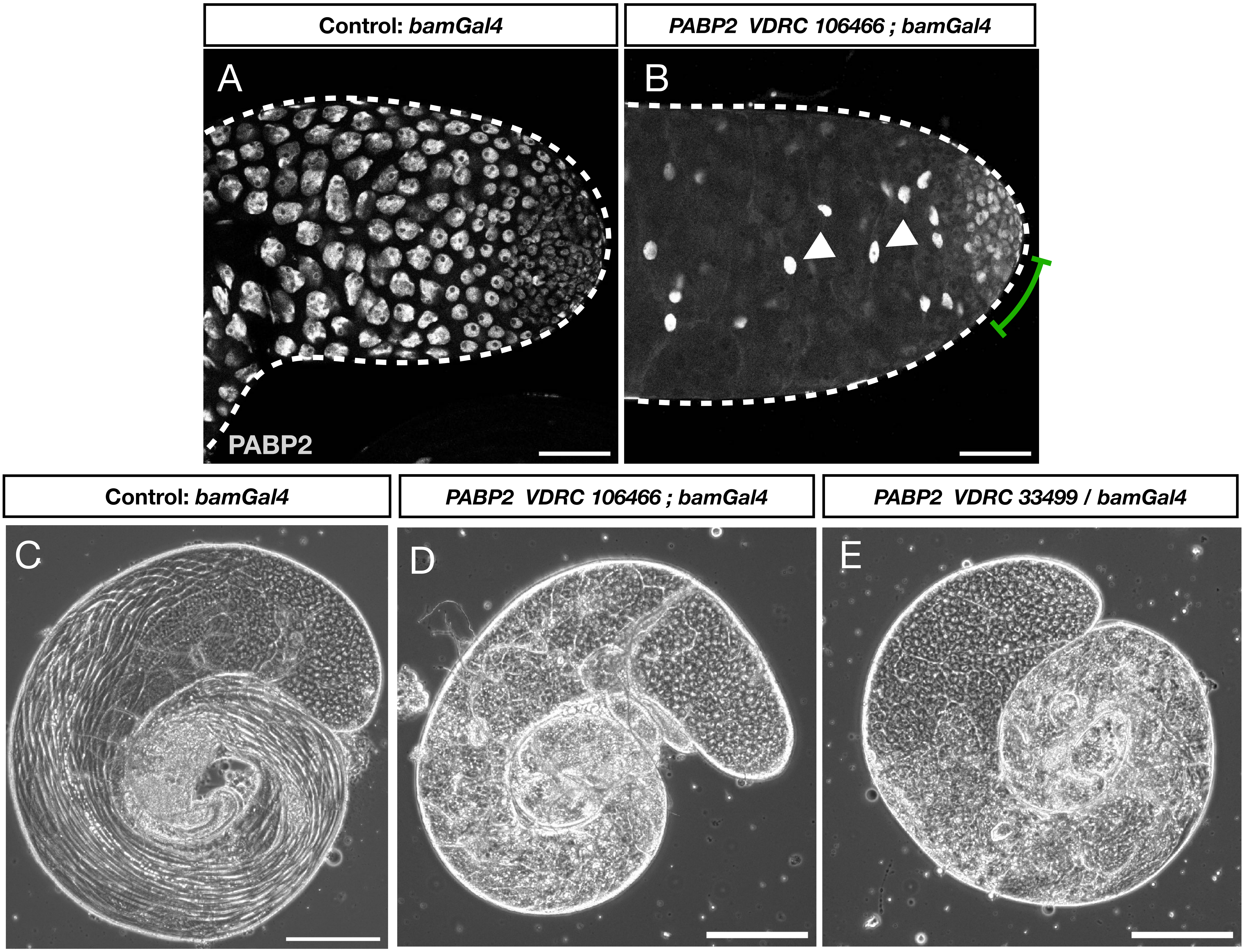
Control: *bamGal4*
*PABP2 VDRC 106466 ; bamGal4*
A
B
PABP2
Control: *bamGal4*
*PABP2 VDRC 106466 ; bamGal4*
*PABP2 VDRC 33499 / bamGal4*
C
D
E

### SOM Figure 5

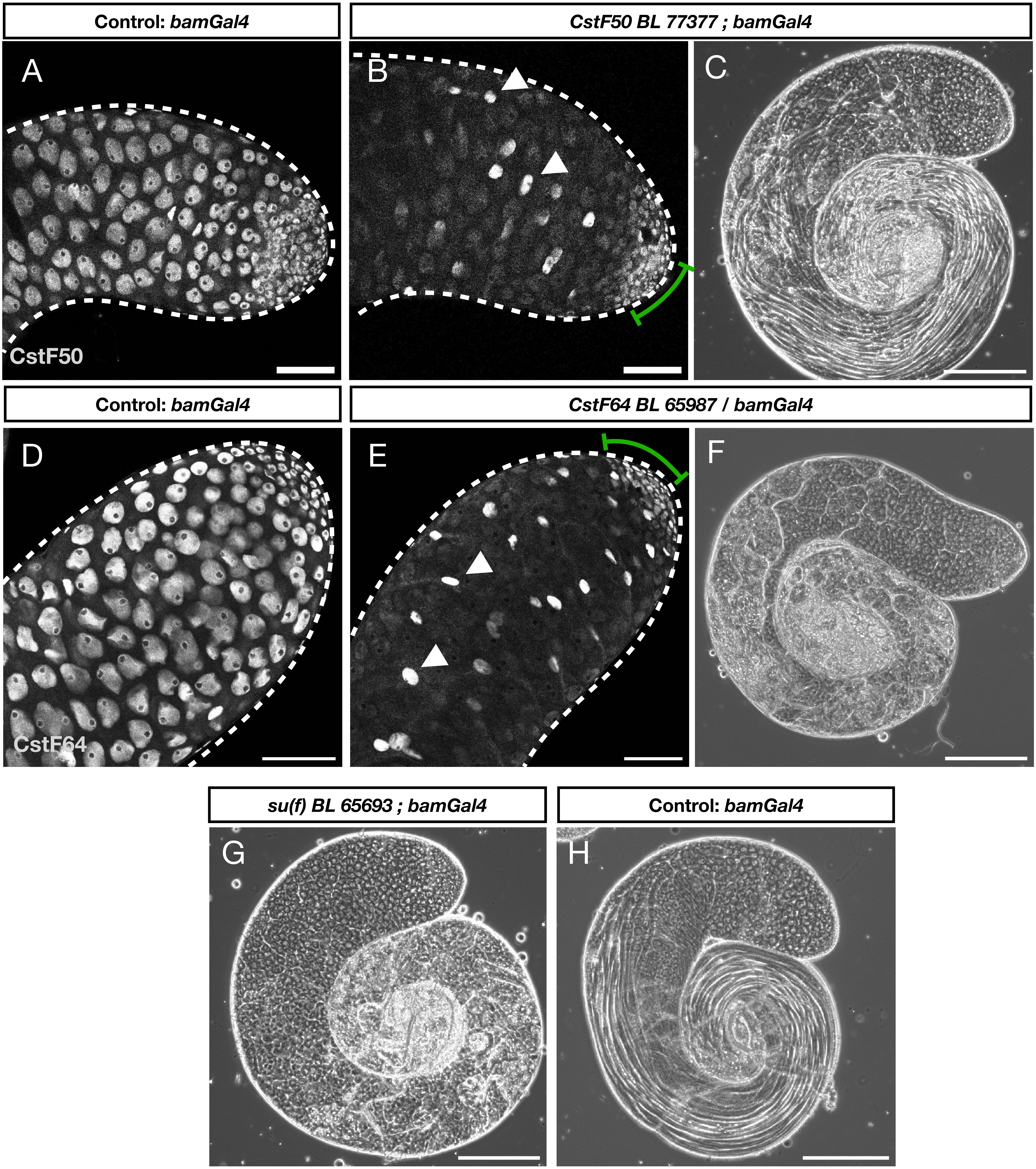
Control: bamGal4
CstF50 BL 77377 ; bamGal4
A
B
C
CstF50
Control: bamGal4
CstF64 BL 65987 / bamGal4
D
E
F
CstF64
su(f) BL 65693 ; bamGal4
Control: bamGal4
G
H

#### SOM Figure 6

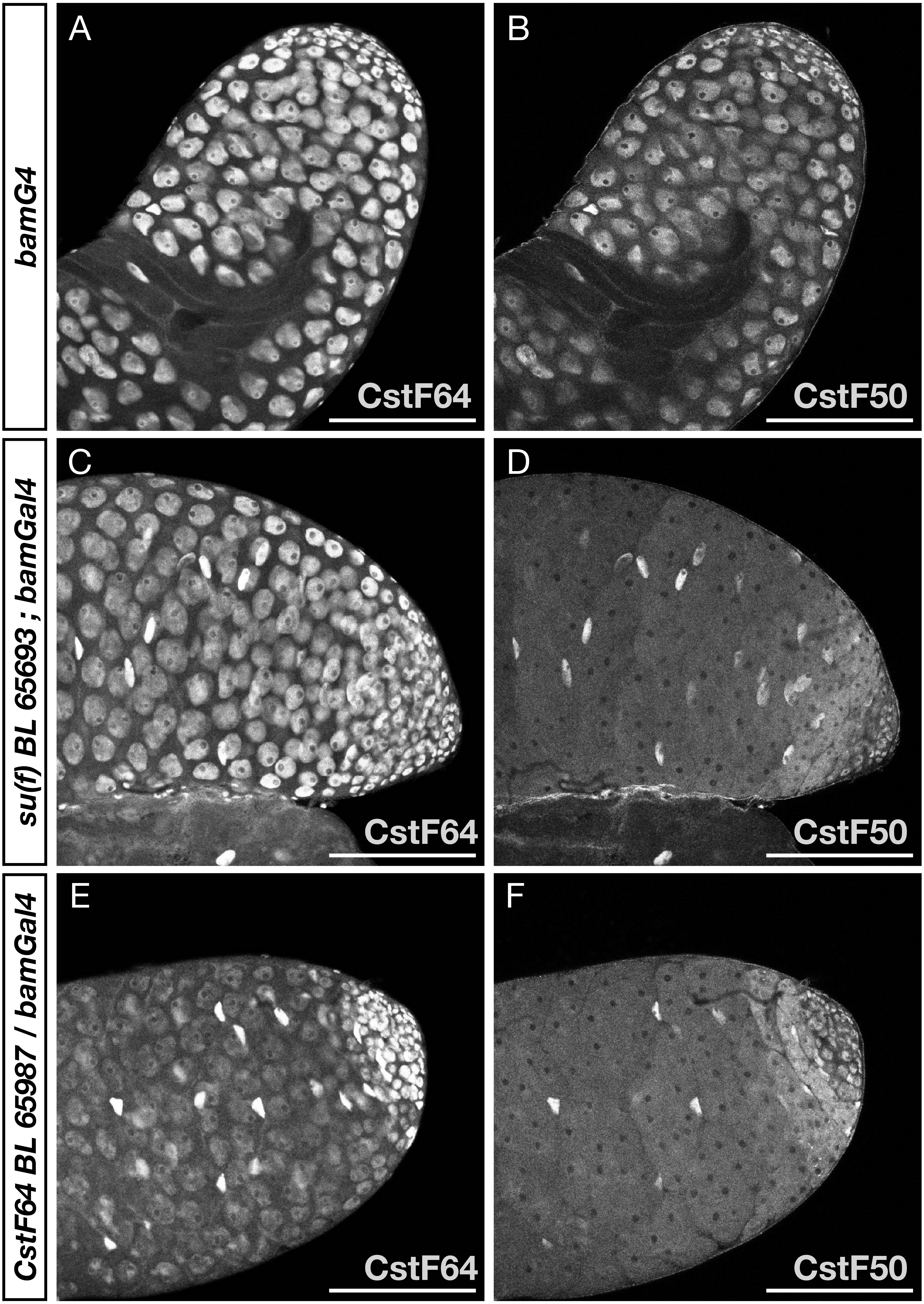

bamG4
su(f) BL 65693 ; bamGal4
CstF64 BL 65987 / bamGal4
A
B
C
D
E
F
CstF64
CstF50
CstF64
CstF50
CstF64
CstF50

### SOM Figure 7

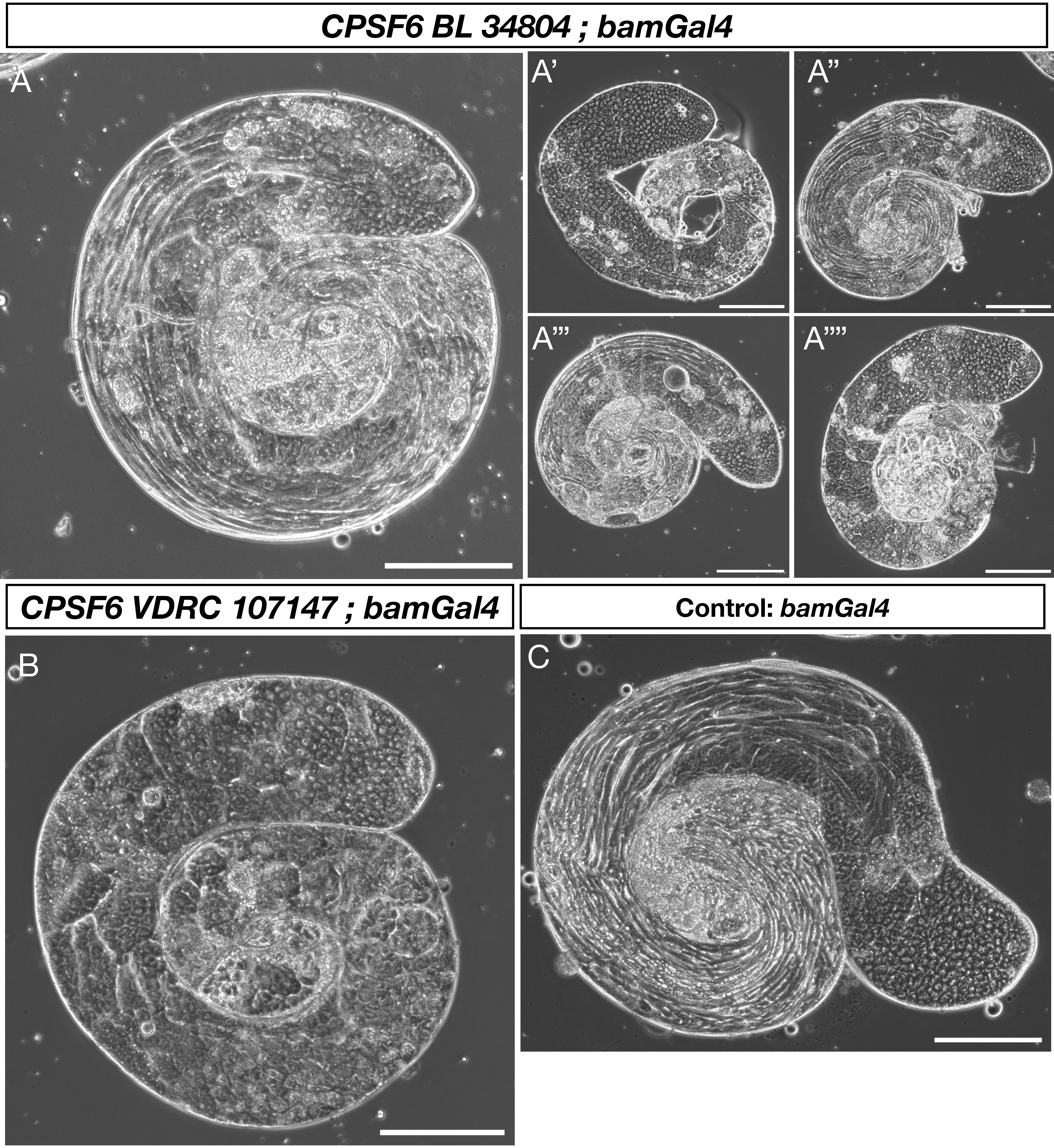
CPSF6 BL 34804 ; bamGal4
A
A’
A”
A’’’
A’’’’
CPSF6 VDRC 107147 ; bamGal4
Control: bamGal4
B
C

### SOM Figure 8

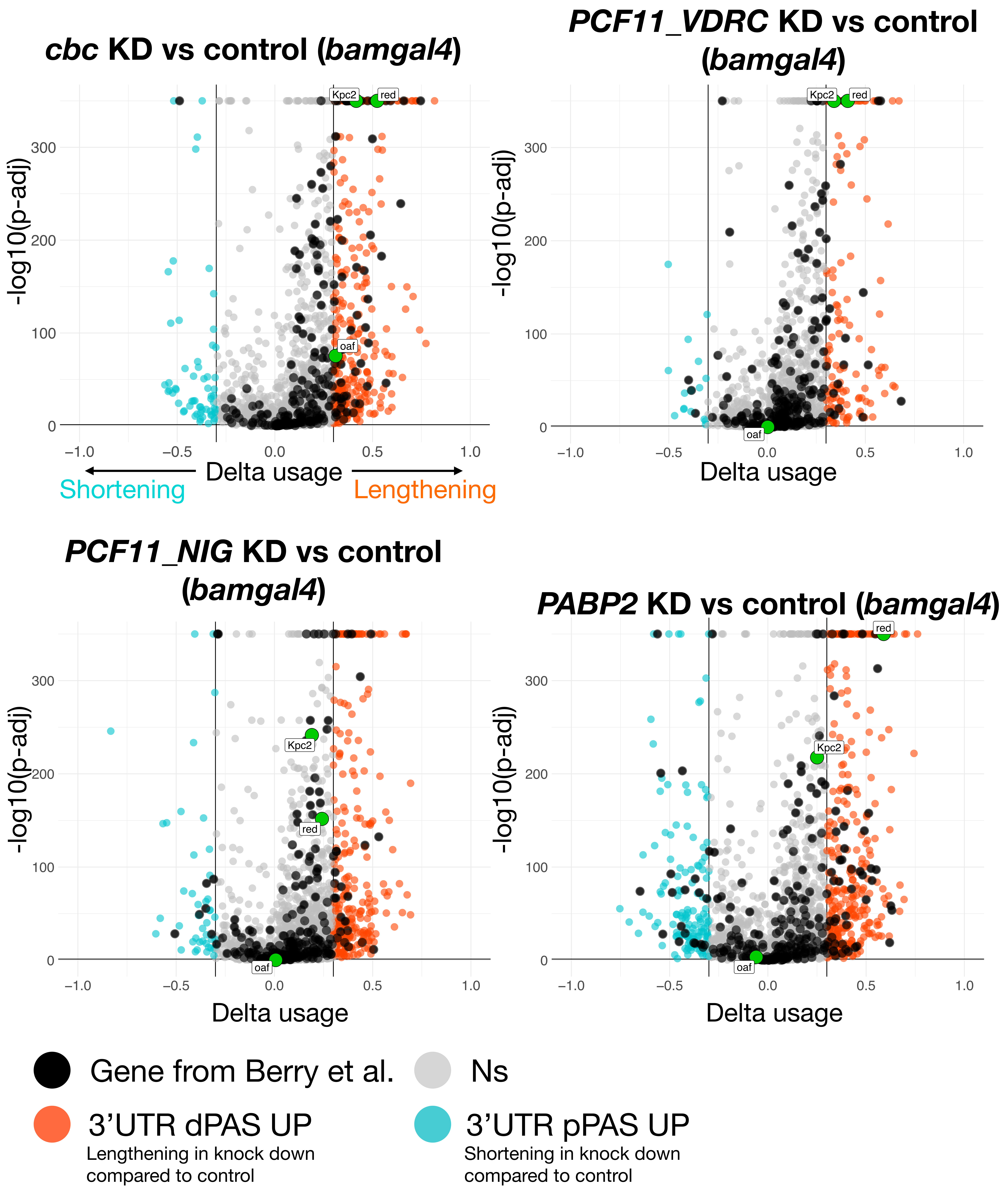

*cbc* KD vs control (*bamgal4*)
*PCF11_VDRC* KD vs control (*bamgal4*)
*PCF11_NIG* KD vs control (*bamgal4*)
*PABP2* KD vs control (*bamgal4*)
-log10(p-adj)
Delta usage
Shortening
Lengthening
Kpc2
red
oaf
Gene from Berry et al.
Ns
3'UTR dPAS UP
Lengthening in knock down compared to control
3'UTR pPAS UP
Shortening in knock down compared to control

### SOM Figure 9

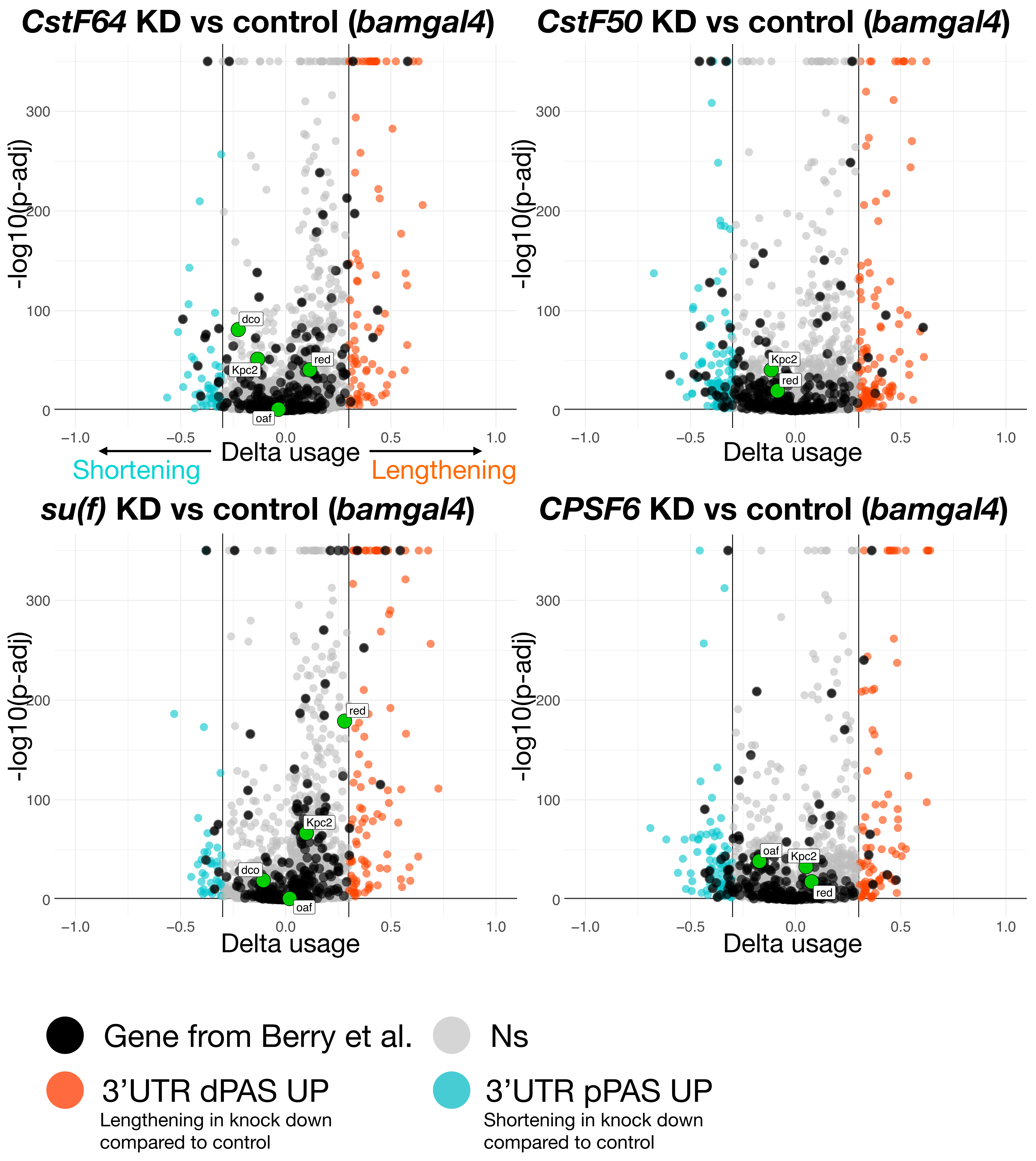

CstF64 KD vs control (bamgal4)
CstF50 KD vs control (bamgal4)
su(f) KD vs control (bamgal4)
CPSF6 KD vs control (bamgal4)
-log10(p-adj)
Delta usage
Shortening
Lengthening
dco
Kpc2
red
oaf
Gene from Berry et al.
Ns
3'UTR dPAS UP
Lengthening in knock down compared to control
3'UTR pPAS UP
Shortening in knock down compared to control

### SOM Figure 10

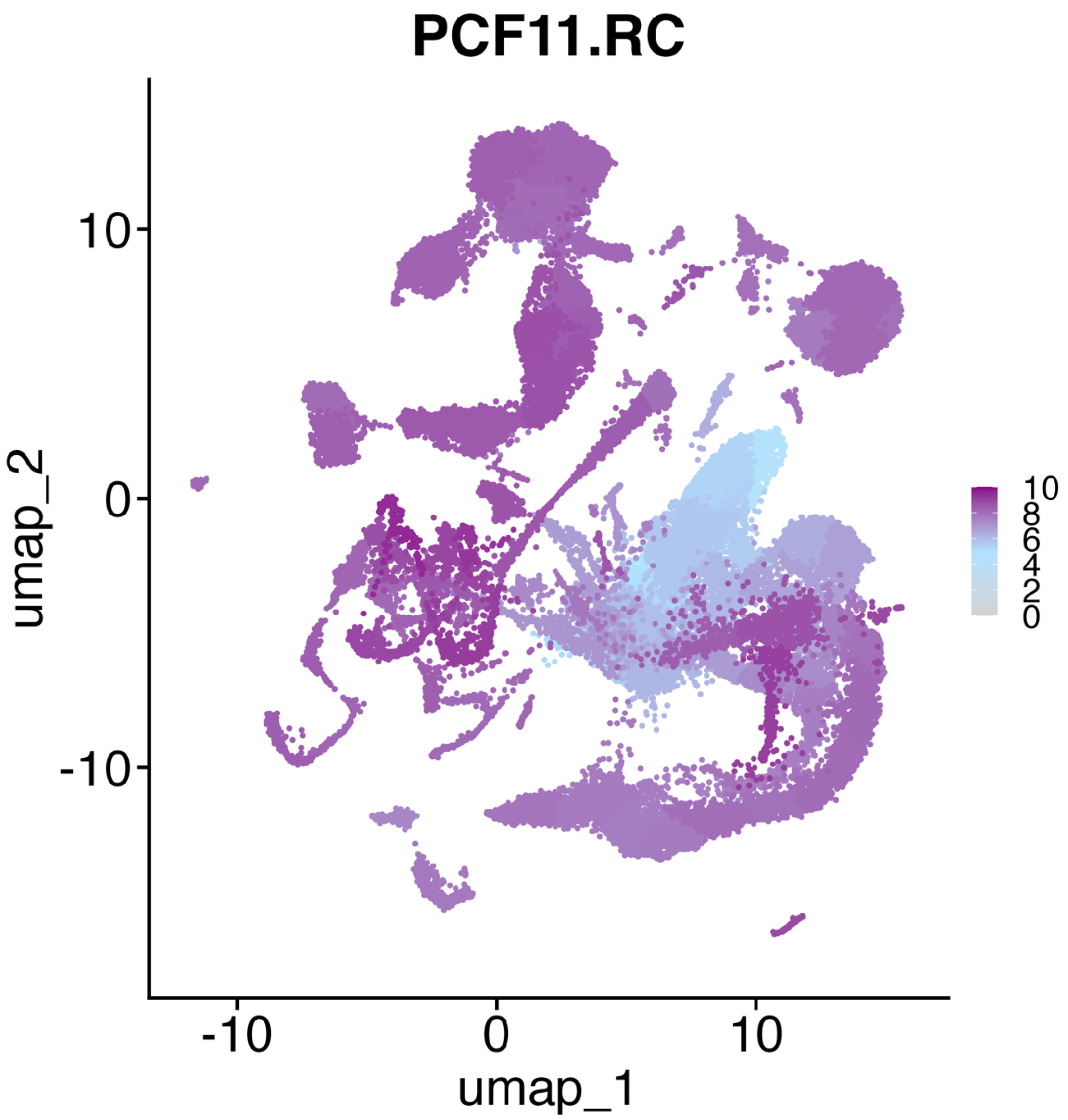

### SOM Figure 11

Western Blot, Fig.5 A-B

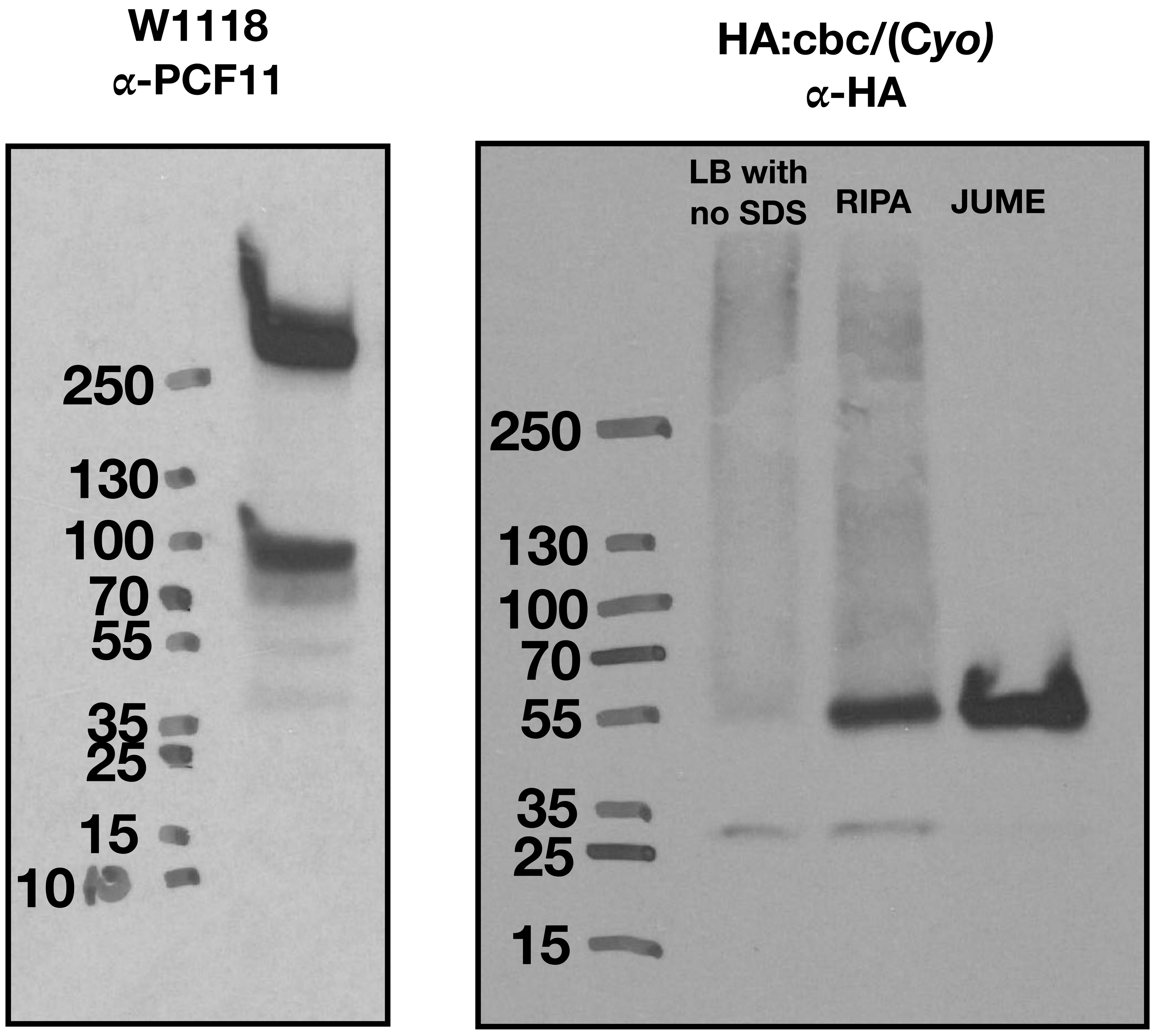


co-IP, Fig.5 C

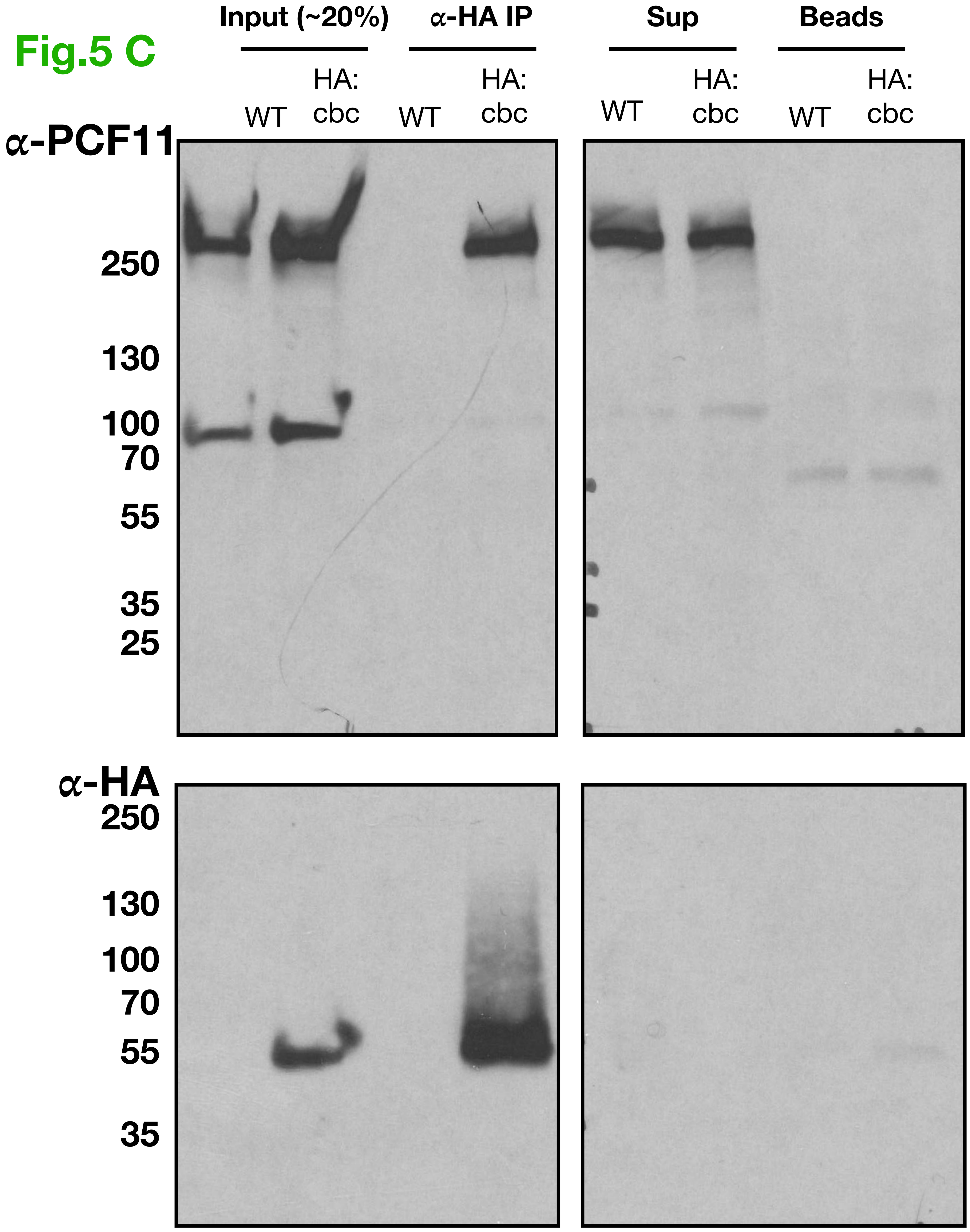

### SOM Figure 12

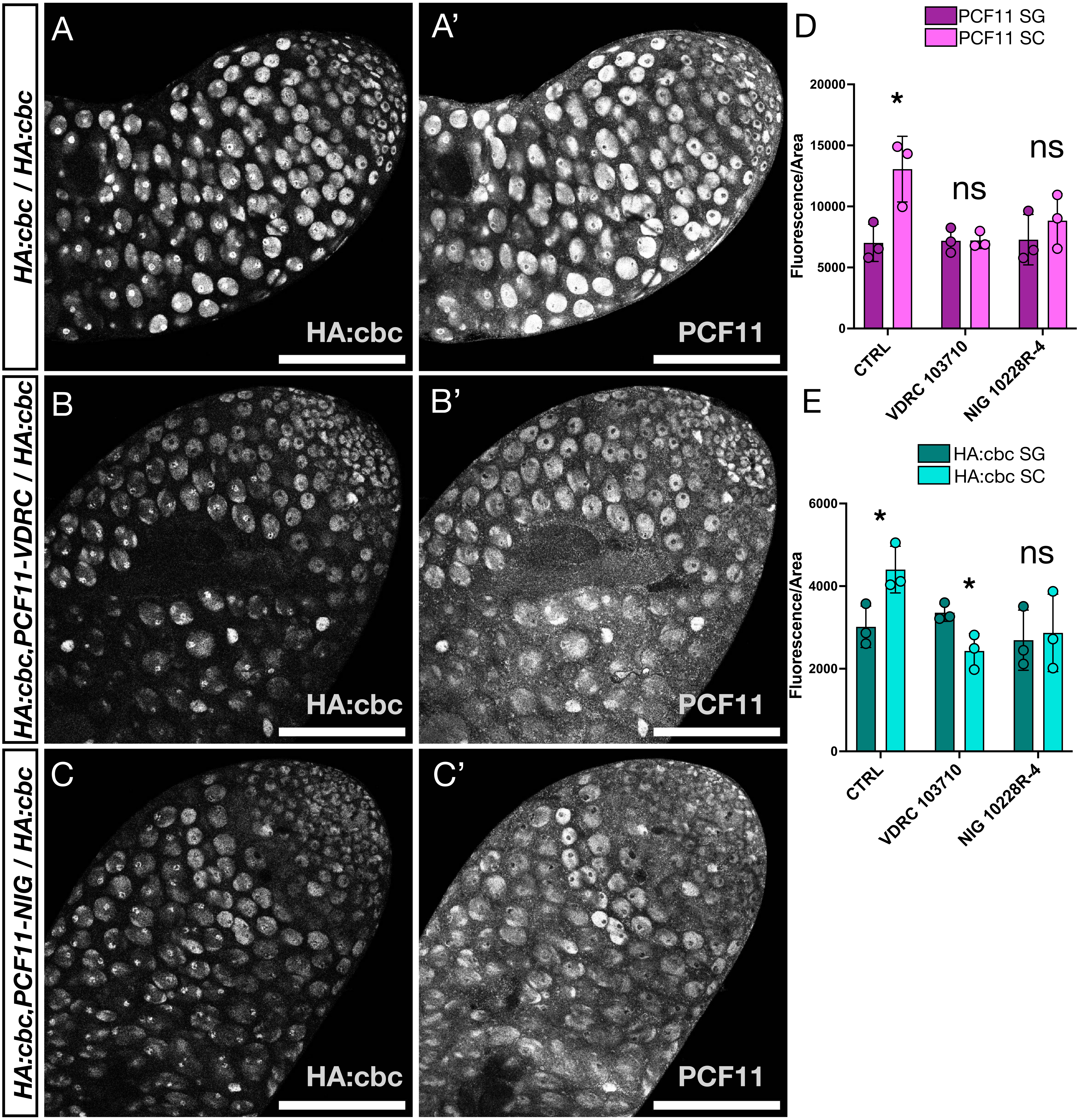

### SOM Figure 13

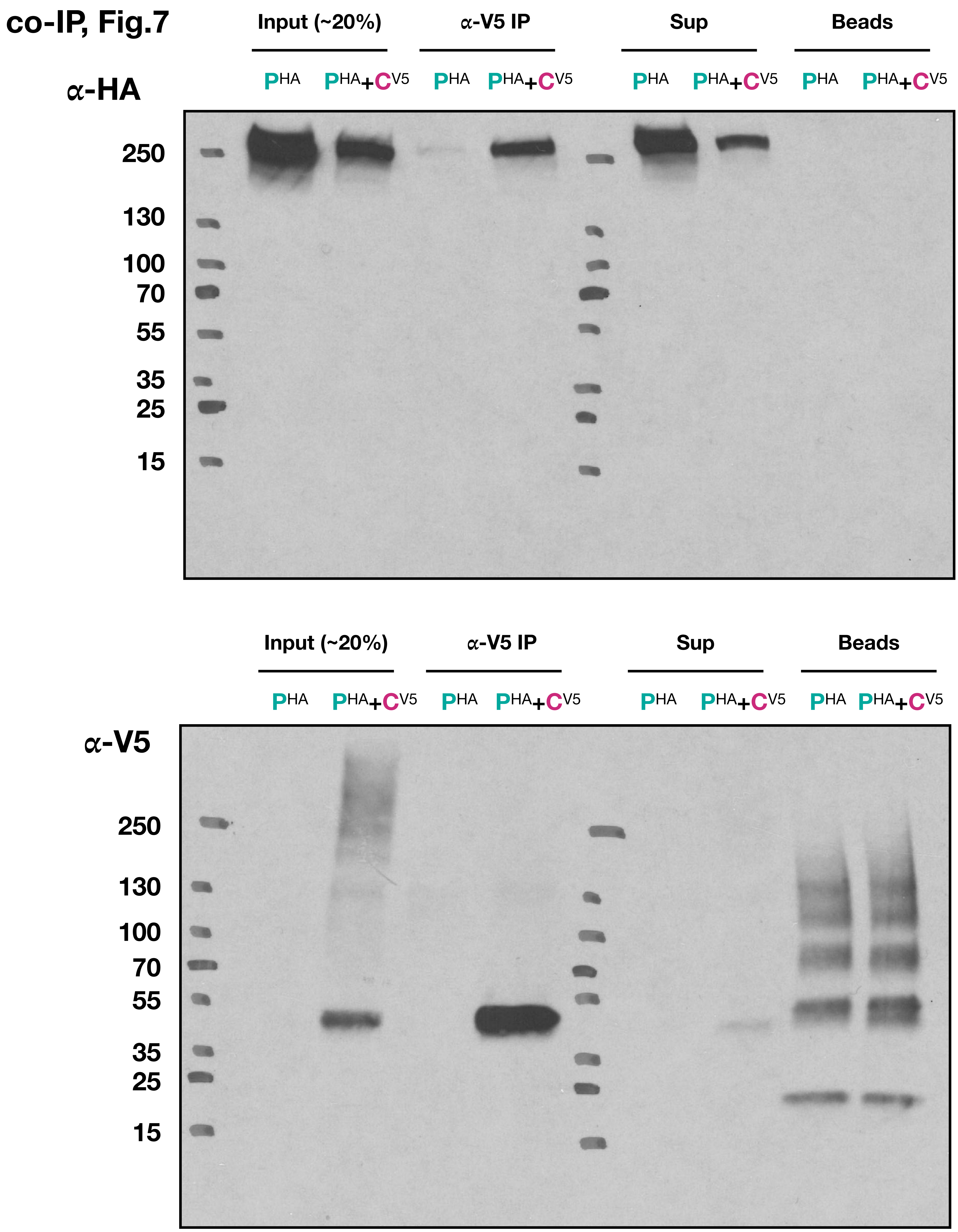

### SOM Figure 14

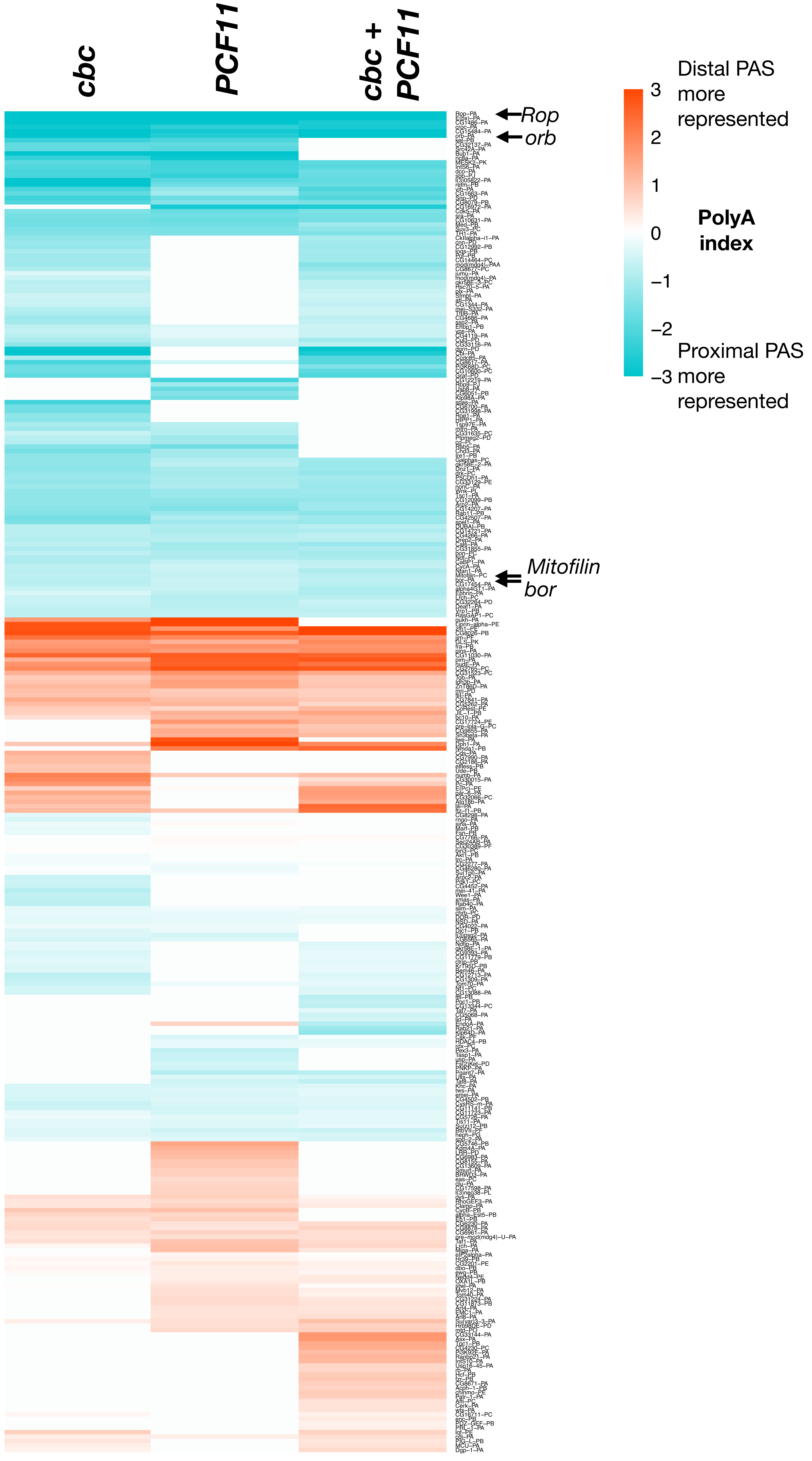

### SOM Figure 15

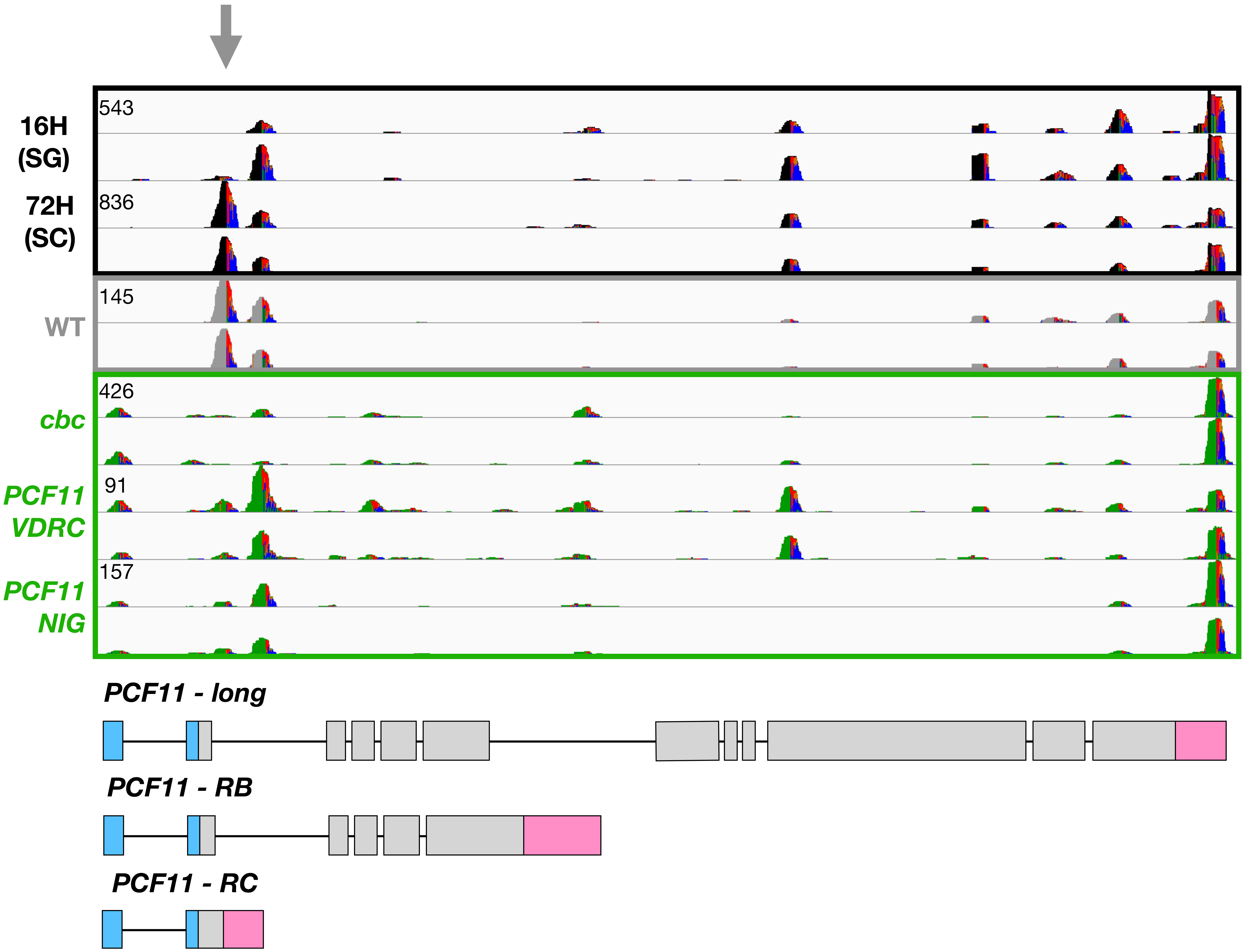
